# GluD2- and Cbln1-mediated Competitive Synaptogenesis Shapes the Dendritic Arbors of Cerebellar Purkinje Cells

**DOI:** 10.1101/2020.06.14.151258

**Authors:** Yukari H. Takeo, S. Andrew Shuster, Linnie Jiang, Miley Hu, David J. Luginbuhl, Thomas Rülicke, Ximena Contreras, Simon Hippenmeyer, Mark J. Wagner, Surya Ganguli, Liqun Luo

**Author notes:** These authors contributed equally (Yukari H. Takeo, S. Andrew Shuster).

## Abstract

The synaptotrophic hypothesis posits that synapse formation stabilizes dendritic branches, yet this hypothesis has not been causally tested *in vivo* in the mammalian brain. Presynaptic ligand cerebellin-1 (Cbln1) and postsynaptic receptor GluD2 mediate synaptogenesis between granule cells and Purkinje cells in the molecular layer of the cerebellar cortex. Here we show that sparse but not global knockout of *GluD2* causes under-elaboration of Purkinje cell dendrites in the deep molecular layer and overelaboration in the superficial molecular layer. Developmental, overexpression, structure-function, and genetic epistasis analyses indicate that dendrite morphogenesis defects result from competitive synaptogenesis in a Cbln1/GluD2-dependent manner. A generative model of dendritic growth based on competitive synaptogenesis largely recapitulates *GluD2* sparse and global knockout phenotypes. Our results support the synaptotrophic hypothesis at initial stages of dendrite development, suggest a second mode in which cumulative synapse formation inhibits further dendrite growth, and highlight the importance of competition in dendrite morphogenesis.

## INTRODUCTION

Nervous system function requires proper dendrite morphogenesis and synapse formation, both of which critically affect neuronal connectivity and integration of synaptic inputs. Disruptions in dendrite morphogenesis and synapse formation are associated with many neurodevelopmental and psychiatric disorders such as autism and schizophrenia (Kulkarni and Firestein, 2012). Research in the past decades has elucidated mechanisms underlying dendrite morphogenesis (Jan and Jan, 2010) and synapse formation (Südhof, 2018; Yogev and Shen, 2014). However, the relationship between these two developmental events is less explored. Dendrite growth is usually considered to occur prior to synapse formation during neuronal differentiation, as a neuron must extend dendritic branches before it can form synapses with incoming axons. However, these two events occur largely concurrently in many cases. Thus, synapse formation and maturation could in principle influence dendritic branching and elongation.

Observations of fixed central nervous system tissue led to the synaptotrophic hypothesis of dendritic growth, which postulates that synapse formation and maturation promote dendritic growth (Cline and Haas, 2008; Vaughn, 1989). Live imaging experiments in amphibians have provided evidence supporting this hypothesis: synapse formation stabilizes nascent dendritic branches, allowing them to stably maintain and extend further (Chen et al., 2010; Niell et al., 2004). A direct test of this hypothesis requires *in vivo* manipulation of synapse formation and visualization of dendritic arbors with single cell resolution. To our knowledge, no such experiments have been performed in the mammalian brain. Here, we investigate this relationship by examining the role of glutamate receptor delta 2 (GluD2)—one of the best-characterized synaptogenic proteins—in dendrite morphogenesis.

GluD2, although belonging to the ionotropic glutamate receptor family, is atypical in that it neither binds glutamate nor exhibits direct channel activity upon binding of known ligands. However, GluD2 has a well-established role in synapse formation and maintenance, as *GluD2* knockout mice lose nearly half of all synapses between the axons of cerebellar granule cells (parallel fibers) and the dendrites of Purkinje cells at the ultrastructural level (Ichikawa et al., 2016; Kashiwabuchi et al., 1995), concomitant with a reduction in physiological responses of Purkinje cells to parallel fiber activation (Kurihara et al., 1997). At these parallel fiber→Purkinje cell synapses, GluD2 present on Purkinje cell dendrites acts as a receptor for cerebellin-1 (Cbln1) secreted by parallel fibers. Cbln1 also binds neurexin, a presynaptic plasma membrane protein on parallel fibers (Yuzaki, 2018). Thus, neurexin, Cbln1, and GluD2 form a tripartite synaptic adhesion complex that promotes parallel fiber→Purkinje cell synapse formation and maintenance (Matsuda et al., 2010; Uemura et al., 2010; Yuzaki, 2018) (**Figure 1A**). In support of the synaptogenic role of this tripartite complex, *Cbln1* knockout also results in up to a ~80% reduction of parallel fiber→Purkinje cell synapses, and addition of recombinant Cbln1 rescues this phenotype in adults (Hirai et al., 2005; Matsuda et al., 2010). Indeed, knockout of *GluD2* or *Cbln1* causes the strongest synapse loss defects among all mouse knockouts of single synaptogenic genes reported. Thus, *GluD2^−/−^* Purkinje cells provide an opportunity to examine the cell-autonomous effects of disrupting synapse formation and maintenance on dendrite growth.

**Figure 1.**
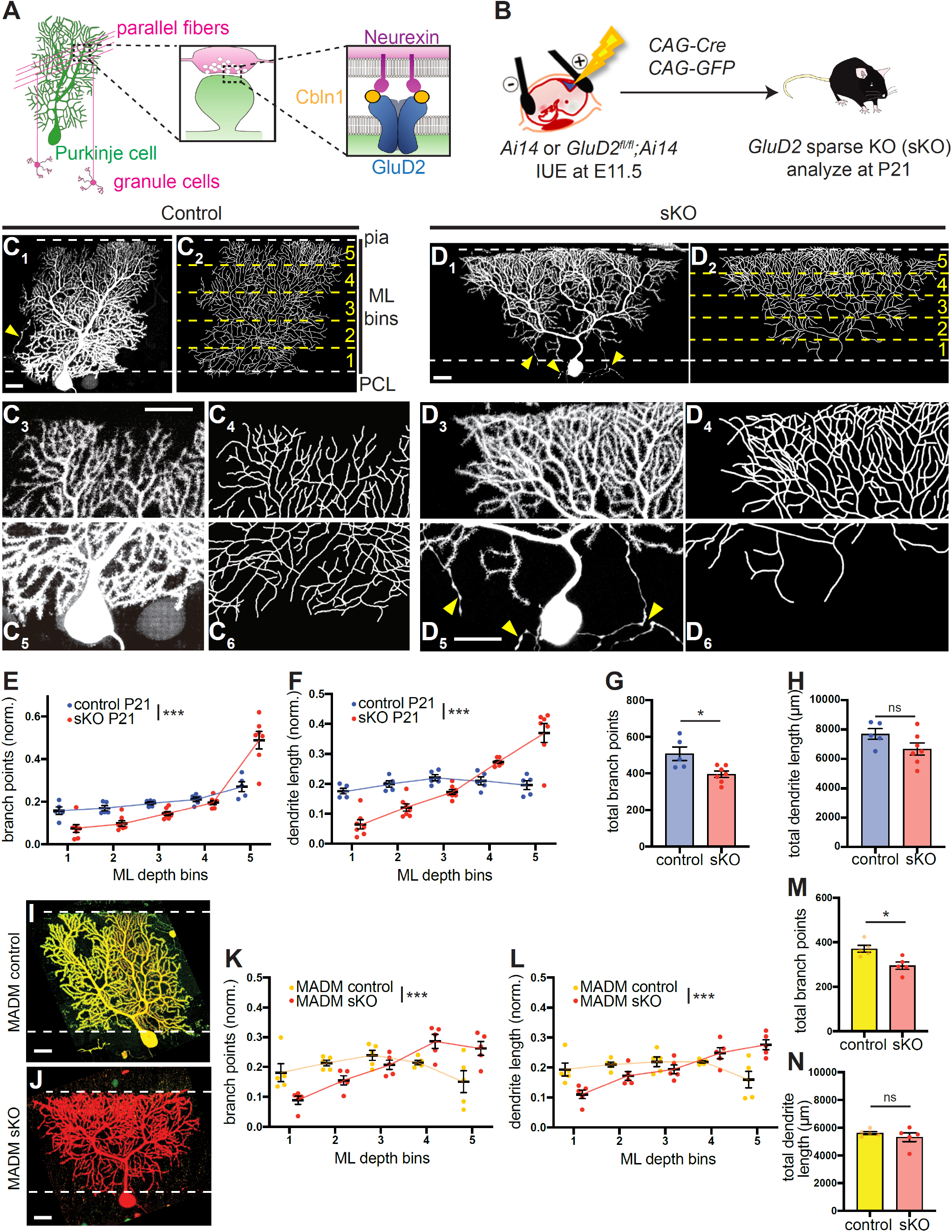
Sparse Knockout of *GluD2* in Purkinje Cells Reduces Dendrite Elaboration in the Deep Molecular Layer but Enhances Dendrite Elaboration in the Superficial Molecular Layer. (A) Schematic summary of parallel fiber→Purkinje cell synaptic connectivity in the cerebellar cortex. Left, anatomy, morphology, and connectivity of cerebellar granule cells (magenta) and Purkinje cells (green). Center, schematic of parallel fiber→Purkinje cell synapses. Right, the tripartite synaptic adhesion complex that promotes parallel fiber→Purkinje cell synapse formation and maintenance. (B) Schematic of *in utero* electroporation (IUE) for genetically accessing Purkinje cells in *GluD2^fl/fl^;Ai14* embryos. Controls were *Ai14* alone. Plasmids encoding Cre recombinase and GFP were co-injected into the fourth ventricle at embryonic day 11.5 (E11.5). Cerebellar samples were collected at postnatal day 21 (P21). (C) Control Purkinje cells labeled via IUE exhibit similar levels of dendritic elaboration in the deep and superficial molecular layer at P21. Representative confocal image of a control Purkinje cell (C_1_) and its tracing (C_2_) are magnified for the superficial (C_3_, C_4_) and deep (C_5_, C_6_) molecular layer. The top and bottom dashed lines in C_1_ and C_2_ represent boundaries between the molecular layer (ML) and the pial surface (pia) and Purkinje cell layer (PCL), respectively. The yellow dashed lines in C_2_ represent divisions between the five numbered bins of the ML. Yellow arrowheads, Purkinje cell axons. Scale bars, 20 μm. (D) Same as (C) except that *GluD2* is knocked out (sKO) of this isolated Purkinje cell via IUE (B). sKO Purkinje cells exhibit an inverted triangular shape, with overelaboration in the superficial molecular layer (D_3_–D_4_) and under-elaboration in the deep molecular layer (D_5_–D_6_). Yellow arrowheads, Purkinje cell axon processes. Scale bars, 20 μm. (E, F) Quantification of the normalized number of branch points (E) and dendrite length (F) in each molecular layer (ML) depth bin in control (blue) and *GluD2* sKO (red) Purkinje cells. Data was normalized to the total number of dendritic branch points or dendrite length across the tree. (G, H) Quantification of the total number of branch points (G) and dendrite length (H) of control and *GluD2* sKO Purkinje cells. Data are mean ± SEM; n = 5 (control), 7 (sKO) cells from 2 (control), 3 (sKO) animals for panels E–H. (I, J) Representative confocal images of a *GluD2^+/−^* (I) and *GluD2^−/−^* (J) cells produced by MADM (see Figure S2A for a scheme). Scale bars, 20 μm. (K, L) Quantification of the normalized number of branch points (K) and dendrite length (L) in each molecular layer (ML) depth bin in MADM control (yellow) and *GluD2* sKO (red) Purkinje cells. (M, N) Quantification of total number of branch points (M) and dendrite length (N) of MADM control and *GluD2* sKO Purkinje cells. Data are mean ± SEM; n = 5 (*GluD2^+/−^* control), 5 (*GluD2^−/−^* sKO) cells from the same animal for panels K–N. Statistics: For binned plots in this (panels E, F, K, L) and all subsequent figures, p-values were calculated using a two-way ANOVA test between control and experimental values in each bin. For bar plots in this (panels G, H, M, N) and all subsequent figures, p-values were calculated using unpaired, two-tailed t tests unless otherwise noted. ns (not significant), p > 0.05; *, p < 0.05; **, p < 0.01; ***, p < 0.001. See Figure S1 and S2 for additional data.

## RESULTS

### Sparse Knockout of *GluD2* in Purkinje Cells Reduces Dendritic Branching in the Deep Molecular Layer but Enhances Dendritic Branching in the Superficial Molecular Layer

To gain genetic access to developing mouse Purkinje cells, we performed *in utero* electroporation (IUE) at embryonic day 11.5 (Nishiyama et al., 2012; Takeo et al., 2015). To delete *GluD2* in Purkinje cells and examine the morphology of mutant dendrites, we electroporated plasmids expressing Cre recombinase and a fluorescent marker into wild-type (control) or *GluD2^fl/fl^* embryos (homozygous for a floxed allele of *GluD2*; Takeuchi et al., 2005) (**Figure 1B**). Typically, only a small fraction of isolated Purkinje cells expressed the plasmids (Nishiyama et al., 2012); therefore, the morphology of the entire dendritic arbor could be easily imaged (**Figure 1C_1_**) and branching patterns completely traced (**Figure 1C_2_**). Likewise, IUE into *GluD2^fl/fl^* embryos resulted in *GluD2* knockout in a sparse population of Purkinje cells in an otherwise wild-type environment; we refer to such cells as sparse knockout (sKO) cells hereafter. Antibody staining validated that Cre/GFP-expressing cells, but not neighboring cells, lacked GluD2 protein (**Figure S1A, B**).

We first examined the morphology of Purkinje cells at postnatal day 21 (P21), when their dendritic arbors have reached the pial surface. In wild-type controls, dendritic arbors, when viewed face-on, typically assumed a nearly rectangular shape in the plane orthogonal to the parallel fibers, with similar widths in the deep (close to Purkinje cell bodies) and superficial (close to the pial surface) portions of the molecular layer (**Figure 1C**). By contrast, *GluD2* sKO Purkinje cells exhibited an inverted triangular shape (**Figure 1D**). In the superficial molecular layer, *GluD2* sKO exhibited much enhanced branching (**Figure 1D_3_, D_4_**) compared to controls (**Figure 1C_3_, C_4_**). Conversely, in the deep molecular layer, *GluD2* sKO Purkinje cells exhibited much reduced arbors (**Figure 1D_5_, D_6_**) compared to controls (**Figure 1C_5_, C_6_**). To quantify these effects, we divided the entire molecular layer into five bins of equal depth and quantified the relative distribution of total dendritic branch points and length within each bin following dendrite tracing (**Figure S1C, D**; **STAR Methods**). Compared to controls, *GluD2* sKO Purkinje cells had fewer branch points and reduced dendritic length in the deep molecular layer but more branch points and increased dendritic length in the most superficial layer (**Figure 1E, F**). When the entire dendritic tree was measured, *GluD2* sKO Purkinje cells exhibited fewer total dendritic branch points but similar total dendritic length as controls (**Figure 1G, H**).

Fine morphological analysis revealed that sKO dendrites had fewer normal-sized dendritic spines and more filopodia-like extensions than control dendrites (**Figure S1E, F**), suggesting that sKO dendrites may persist in an immature, exploratory state compared to controls. This is consistent with the established requirement for GluD2 in synapse formation and maintenance (Kurihara et al., 1997). Previous studies have also shown that in *GluD2* knockout mice, climbing fibers (axons from inferior olive neurons), which normally innervate the deepest 80% of Purkinje cell dendritic trees, invade more superficial depths of the molecular layer (Hirai et al., 2005; Ichikawa et al., 2002). *GluD2* sKO cells exhibited this phenotype when examined at P63 (**Figure S1G**).

To validate the *GluD2* sKO dendrite morphogenesis phenotypes via an independent method, we used mosaic analysis with double markers (MADM; Contreras et al., 2020; Zong et al., 2005) to knock out *GluD2* in a sparse population of neurons uniquely marked in an otherwise *GluD2^+/−^* background (**Figure S2A**). MADM-mediated sKO was also confirmed by a loss of GluD2 protein (**Figure S2B, S2C**). Compared to MADM *GluD2^+/−^* controls (**Figure 1I, Figure S2D**), which resembled wild-type cells, P21 MADM *GluD2^−/−^* Purkinje cells exhibited dendrite morphogenesis defects similar to those generated via IUE-based sKO, with reduced branching in the deep molecular layer and enhanced branching in the superficial molecular layer (**Figure 1J, Figure S2E**). Quantifying the distribution of dendritic branches and lengths in five depth bins across the molecular layer revealed phenotypes in MADM *GluD2* sKO cells (**Figure 1K, L**) similar to IUE-mediated *GluD2* sKO cells (**Figure 1G, H**). Similarly, MADM *GluD2* sKO cells exhibited fewer total dendritic branch points than but similar total dendritic length to controls (**Figure 1M, N**). Together, these findings demonstrate that GluD2, a receptor essential for parallel fiber→Purkinje cell synapse formation, cell-autonomously regulates Purkinje cell dendrite branching.

### Global Knockout of *GluD2* Does Not Recapitulate Purkinje Cell Dendrite Morphogenesis Phenotypes Caused by Sparse Knockout

The *GluD2* sKO phenotypes described above were unexpected, as previous studies of Purkinje cell morphology in germline *GluD2* knockout animals did not report such phenotypes (Kaneko et al., 2011; Kashiwabuchi et al., 1995). To determine if the phenotypes are specific to sparse knockout, we sparsely labeled wild-type control and *GluD2* germline (global) knockout (gKO) Purkinje cells using IUE (**Figure 2A**). Consistent with previous reports, we did not find gross changes in the morphology of *GluD2* gKO Purkinje cell dendritic arbors compared to controls (**Figure 2B, C**; **Figure S2F, G**; quantified in **Figure 2D, E**). The total number of branch points and dendritic length in *GluD2* gKO cells were not significantly different from controls (**Figure 2F, G**). Like sKO cells, gKO cells also had fewer normal-sized spines compared to control cells (**Figure S1E, F**), consistent with GluD2’s essential role in synaptogenesis.

**Figure 2.**
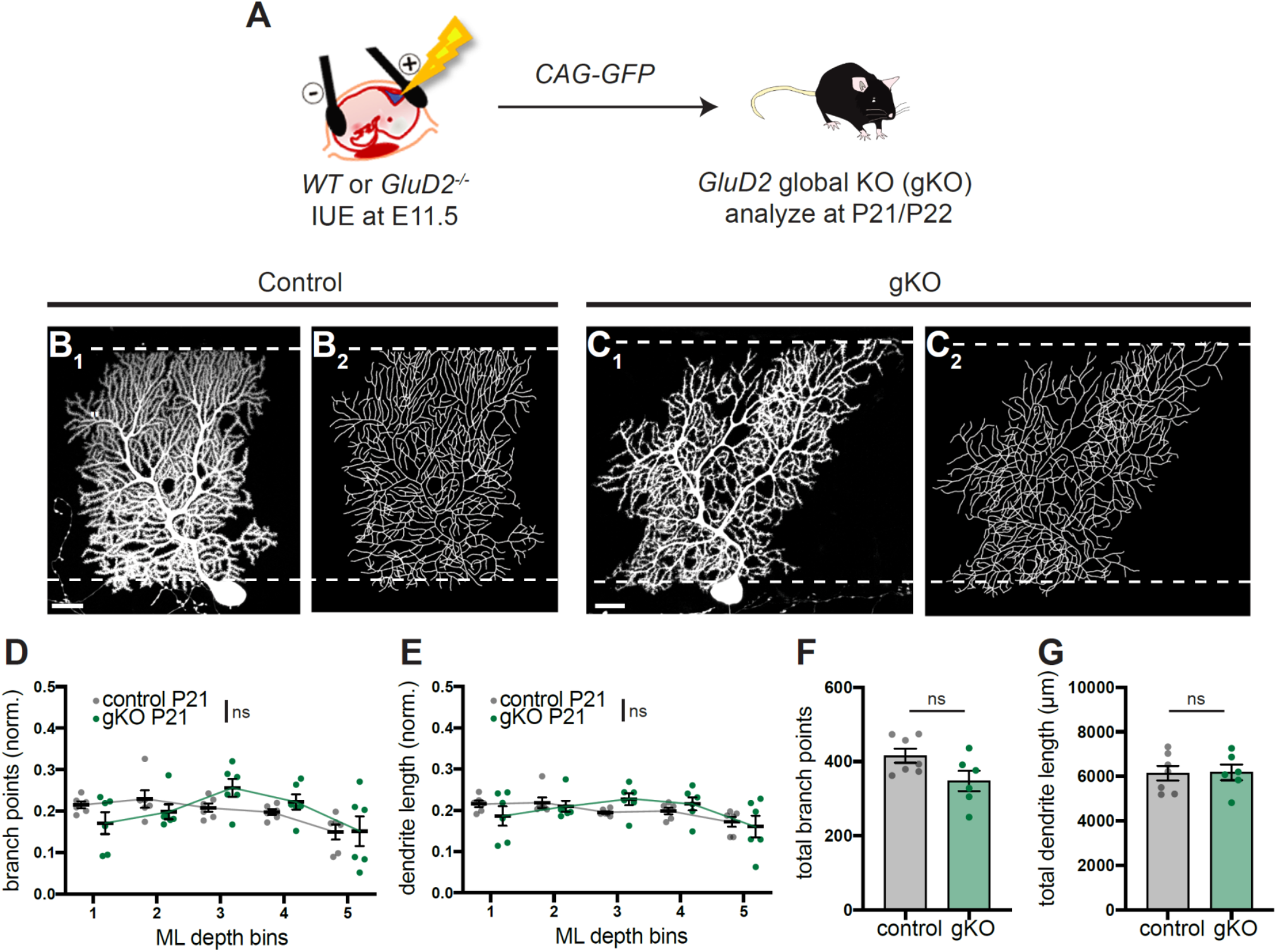
Global Knockout of *GluD2* Does Not Recapitulate Purkinje Dendrite Morphogenesis Phenotypes Caused by Sparse Knockout. (A) Schematic of *in utero* electroporation (IUE) for genetically accessing Purkinje cells in *wild-type* (WT) and *GluD2^−/−^* (gKO) embryos. Plasmids encoding GFP were injected into the fourth ventricle at embryonic day 11.5 (E11.5). Cerebellar samples were collected on postnatal days 21 or 22 (P21/22). (B, C) Wild-type control and global *GluD2^−/−^* (gKO) Purkinje cells both exhibit similar levels of dendritic elaboration in the deep and superficial molecular layer around P21, as seen in representative confocal images (B_1_, C_1_) Purkinje cells and tracings (B_2_, C_2_) of Purkinje cell dendritic arbors. Scale bars, 20 μm. (D, E) Quantification of the normalized number of branch points (D) and dendrite length (E) in each molecular layer (ML) bin in wild-type control (grey) and *GluD2* gKO (green) Purkinje cells. Data are mean ± SEM; n = 6 (control), 6 (gKO) cells from 2 (control), 2 (gKO) animals; ns, not significant (p > 0.05 by two-way ANOVA). (F, G) Quantification of total number of branch points (F) and dendrite length (G) of control (grey) and *GluD2* gKO (green) Purkinje cells. Data are mean ± SEM; n = 6 (control), 6 (gKO) cells from 2 (control), 2 (gKO) animals; p-values were calculated using unpaired t tests. See Figure S2 for related data.

Taken together with the sKO phenotypes, these data revealed that Purkinje cell dendrite morphology is affected more by relative differences in GluD2 signaling between neighboring Purkinje cells than by the absolute level of GluD2 signaling within individual cells. Given the established function of GluD2 in parallel fiber→Purkinje cell synaptogenesis, these data suggest that the GluD2 sparse knockout dendritic branching phenotypes may result from Purkinje cell dendrites competing with their neighbors for access to parallel fibers in the process of forming synapses.

### *GluD2* Sparse Knockout Phenotypes Arise Early During Postnatal Cerebellar Development and Persist in Adults

When does the *GluD2* sKO phenotype arise during development? Cerebellar morphogenesis occurs primarily during the first three postnatal weeks. At birth, granule cell progenitors occupy the most superficial layer (the external granular layer), where they undergo rapid proliferation. As granule cells exit mitosis, they extend their axons as parallel fibers in the space between granule cell progenitors and Purkinje cells, which gradually develop into the molecular layer, while their cell bodies migrate past the Purkinje cell layer to descend into the internal granular layer, giving rise to the granule cell layer in adults (Altman, 1972a) (**Figure 3A**). Later-born granule cells stack their parallel fibers superficially to those from earlier-born granule cells (Espinosa and Luo, 2008). Developing parallel fibers are the substrate on which Purkinje cell dendrites grow, branch, and form synapses, expanding the molecular layer in the process. As granule cell neurogenesis proceeds, the external granular layer is gradually replaced by the molecular layer until P21, when granule cell neurogenesis is complete and Purkinje cell dendrites reach the pial surface (**Figure 3A**).

**Figure 3.**
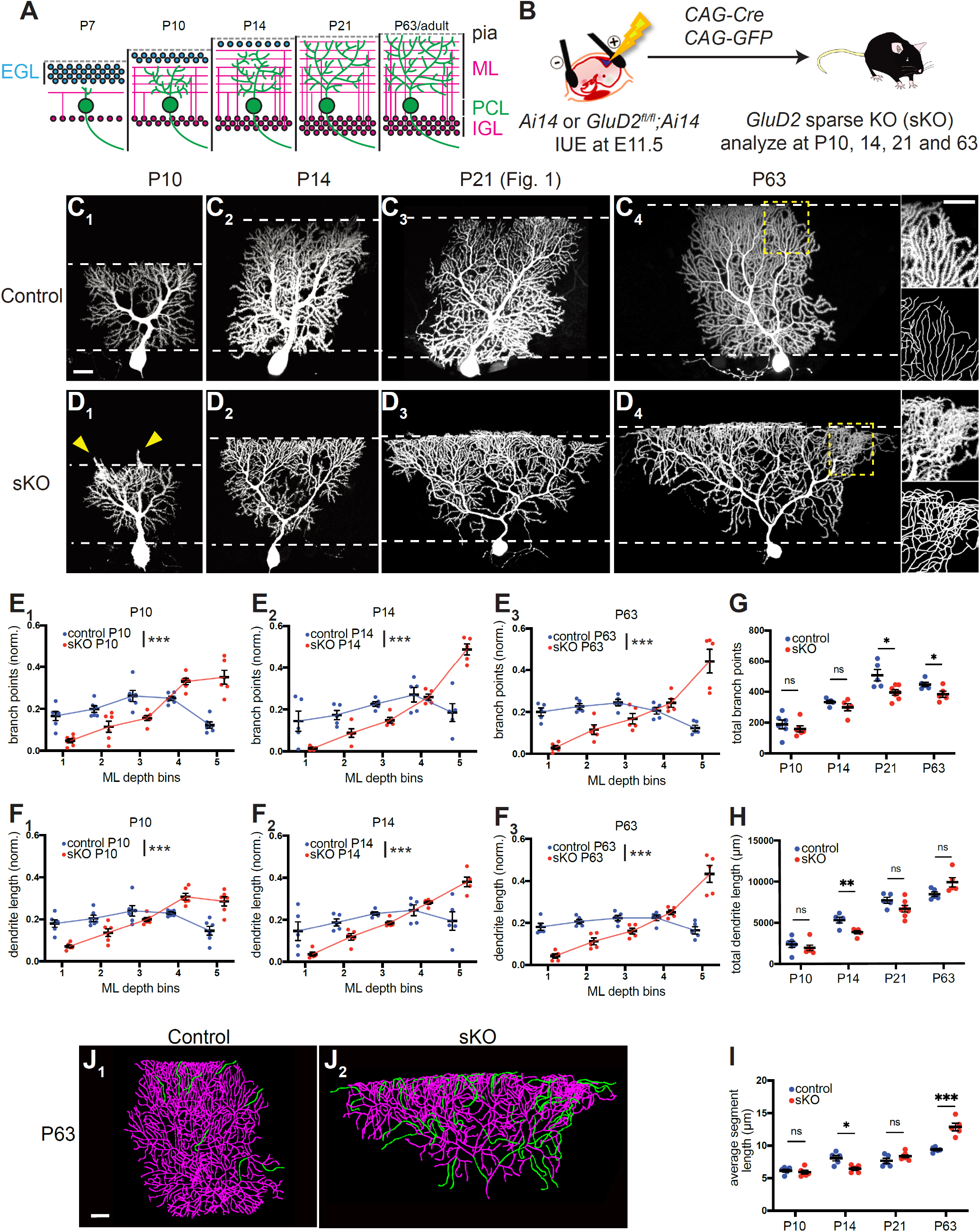
*GluD2* Sparse Knockout Phenotypes Arise Early During Postnatal Cerebellar Development and Persist into Adulthood. (A) Schematic of the time course of postnatal Purkinje cell dendritic morphogenesis. Pia, pial surface; EGL, external germinal layer; ML, molecular layer; PCL, Purkinje cell layer; IGL, internal granular layer. (B) Schematic of *in utero* electroporation (IUE) for genetically accessing Purkinje cells in *Ai14* and *GluD2^fl/fl^;Ai14* embryos. (C, D) Control Purkinje cells (C) exhibit similar levels of dendritic elaboration in the deep and superficial molecular layer between P10 and adulthood (P63). *GluD2* sKO Purkinje cells (D) exhibit an inverted triangular shape, with under-elaboration of dendrites in the deep molecular layer and overelaboration of dendrites in the superficial molecular layer beginning around P10 and persisting into adulthood (P63). Arrowheads in D_1_ indicate overextension of dendritic branches beyond the superficial border of the molecular layer. Higher magnification of superficial portions of dendritic trees and traced images are shown in insets in C_4_ and D_4_, highlighting frequent dendritic branch crossing-over in sKO, but not control, cells. Scale bar, 20 μm. (E, F) Quantification of the normalized number of branch points (E) and dendrite length (F) in each molecular layer (ML) bin in control (blue) and *GluD2* sKO (red) Purkinje cells at P10, P14, and P63 (see Figure 1E, F for analogous quantifications at P21). (G–I) Quantification of the total number of branch points (G), dendrite lengths (H) and average segment lengths (I) of control (blue) and *GluD2* sKO (red) Purkinje cells. For panel E–I, data are mean ± SEM; n = 6 (P10 control), 6 (P10 sKO), 5 (P14 control), 5 (P14 sKO), 5 (P63 control), 5 (P63 sKO) cells from 2 (P10 control), 2 (P10 sKO), 2 (P14 control), 2 (P14 sKO), 2 (P63 control), 4 (P63 sKO) animals; p-values were calculated two-way ANOVA between control and sKO values in each bin (for E, F) or unpaired t-tests (for G–I). P21 data are the same as those in Figure 1. (J) Representative tracings (from C_4_ and D_4_) of control and *GluD2* sKO Purkinje cell dendritic arbors with dendritic segments longer than 30 μm highlighted in green. Scale bar, 20 μm. See Figure S3 for related data.

To characterize the role GluD2 plays in dendrite morphogenesis across development, we profiled the developmental trajectory of the *GluD2* sKO phenotype by analyzing dendritic arbors of Purkinje cells at postnatal days 7, 10, 14, 21, and ~63 using IUE (**Figure 3B**). We found reduced branching in the deep molecular layer and over-branching in the superficial molecular layer of *GluD2* sKO cells as early as P10 (**Figure 3C_1_, D_1_**; see P7 analysis in **Figure S3A–I**), when parallel fibers of granule cells are already forming synapses onto Purkinje cells (Altman, 1972b; West and del Cerro, 1976). These phenotypes persisted across all later stages, with the superficial molecular layer over-branching phenotype becoming more pronounced with age (**Figure 3C_2_–C_4_, D_2_–D_4_**; see additional images in **Figure S3J–L**). Quantification of distributions of dendritic branch points (**Figure 3E_1_–E_3_**; **Figure 1E**) and length (**Figure 3F_1_–F_3_**; **Figure 1F**) supported the above observations. We note that at P10, *GluD2* sKO cells had increased occurrence of dendritic branches extending beyond the superficial border of the molecular layer (arrowheads in **Figure 3D_1_** and **Figure S3J_1_**, quantified in **Figure S3J_2_**). Quantification of total dendritic branch points, lengths, and average segment lengths (**Figure 3G–I**) indicated a general trend of fewer total dendritic branch points in sKO cells, reaching statistical significance at P21 and P63 (**Figure 3G**). A trend of decreased total dendritic length in sKO cells appeared in early stages but was reversed at P63 (**Figure 3H**). The combination of the above resulted in a highly significant increase in the average dendritic segment length of sKO dendrites compared to controls at P63 (**Figure 3I**), mostly due to long terminal branches (**Figure 3J, S4J**). In addition, P21 and particularly P63 sKO Purkinje dendritic trees exhibited crossing-over of dendritic branches caused by branches that extended out of the 2D plane that normal dendritic trees are restricted to (compare **Figure 1D_3_, D_4_** with **Figure 1C_3_, C_4_**; insets of **Figure 3D_4_** with **3C_4_**).

In summary, our developmental analyses revealed that GluD2 sKO cells exhibited reduced branching and elongation in the deep molecular layer accompanied by increased dendritic branching and elongation in the superficial molecular layer at all ages examined. Together, these two phenotypes caused the inverted triangular dendrite morphology we observed.

### GluD2 Overexpression Causes Dendritic Over-branching in the Deep Molecular Layer

To complement the sparse knockout studies, we next examined the effects of sparsely overexpressing GluD2 (GluD2-OE) in Purkinje cells of wild-type mice via IUE (**Figure 4A**). At P7, regions adjacent to GluD2-OE Purkinje cell dendrites exhibited increased staining intensity of the vesicular glutamate transporter vGluT1, a presynaptic marker only expressed by granule cells in the cerebellar cortex (Miyazaki et al., 2003; **Figure 4B, C**), consistent with the synaptogenic role of GluD2. vGluT1 staining levels positively correlated with GluD2-OE levels (**Figure 4D**). Compared with controls (**Figure 4E**), GluD2-OE Purkinje cells also exhibited two additional phenotypes at P7: supernumerary primary dendrites (**Figure 4F, G**) and numerous spine-like protrusions from dendritic segments (**Figure 4F_2_**), which were largely absent on control Purkinje cells at this stage (**Figure 4E_2_**). To quantify these effects, we categorized imaged Purkinje cells as having 1–2 primary dendrites or 3+ primary dendrites and, independently, as “spiny” or “non-spiny”, blind to genotype and GluD2 level. 81% of wild-type controls at P7 had 1–2 primary dendrites, whereas only 35% of GluD2-OE cells had 1–2 primary dendrites (**Figure 4G**). Furthermore, whereas all control cells were non-spiny, 92% of GluD2-OE cells with 3+ primary dendrites were spiny (**Figure 4H**). The minority of non-spiny GluD2-OE cells tended to have lower GluD2 levels than the more typical spiny cells (**Figure 4D**). These results suggest that GluD2 overexpression promotes Purkinje cell synaptogenesis, with accompanying exuberant dendritic branching.

**Figure 4.**
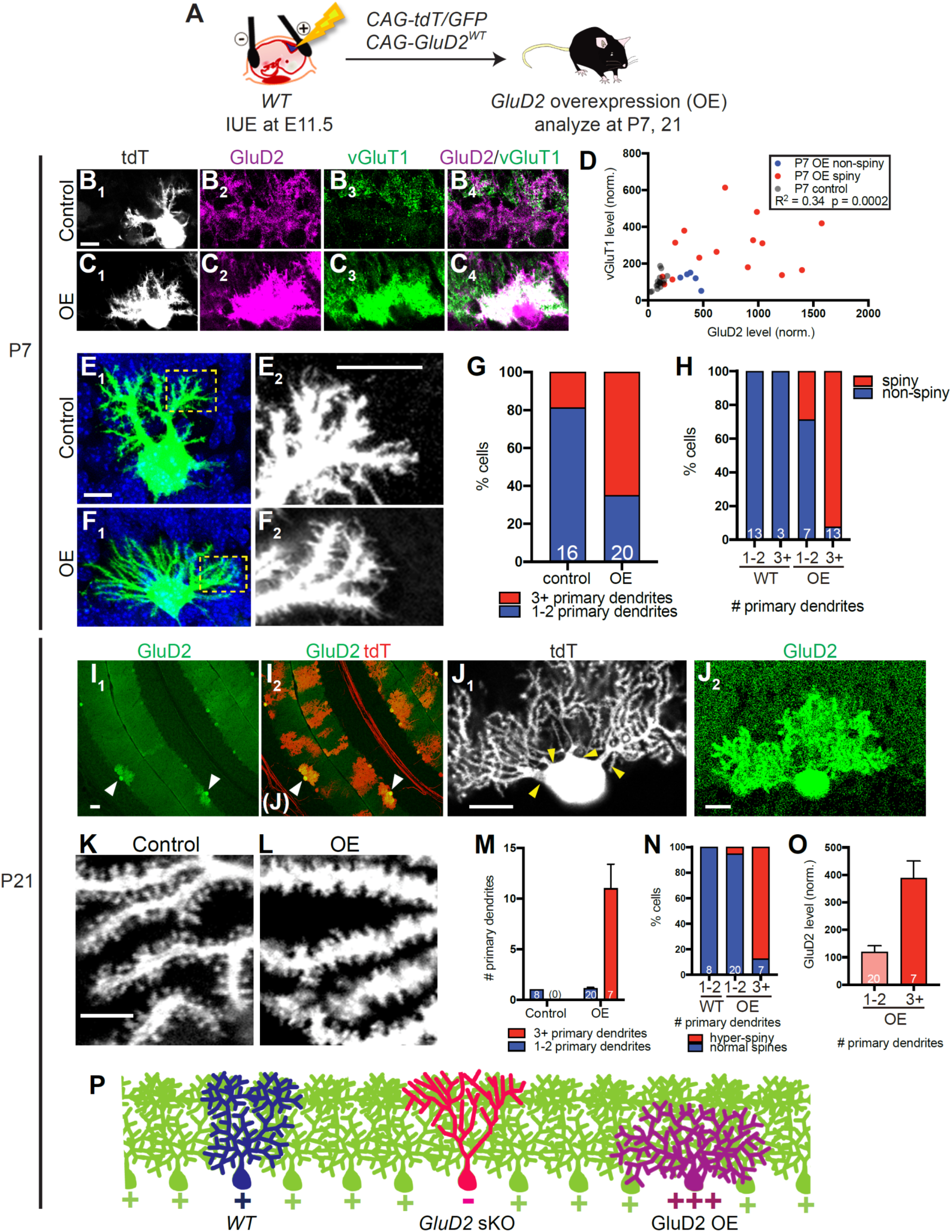
GluD2 Overexpression Causes Dendrite Overelaboration in the Deep Molecular Layer. (A) Schematic of *in utero* electroporation (IUE) for overexpressing GluD2 in Purkinje cells (GluD2-OE) of wild-type embryos. Plasmids encoding tdT/GFP and GluD2^WT^ were co-injected into the fourth ventricle at embryonic day 11.5 (E11.5). Controls did not receive *CAG-GluD2^WT^* plasmid. Cerebellar samples were collected at postnatal days 7 (P7) and 21. (B, C) tdTomato expression (B_1_, C_1_), immunostaining for GluD2 (B_2_, C_2_) and vGluT1 (B_3_, C_3_) and merge (B_4_, C_4_) in control (B) and GluD2-OE (C) Purkinje cells at P7. Scale bar, 10 μm. (D) Plot of the relationship between vGluT1 and GluD2 levels of each P7 control and GluD2-OE cell (dot). (E, F) Representative images of control (E_1_) and GluD2-OE (F_1_) Purkinje cells at P7, showing supernumerary primary dendrites on the GluD2-OE cell. Green, tdT in Purkinje cell. Blue, DAPI. E_2_ and F_2_ are high magnification images of distal dendrites of E_1_ and F_1_ (boxed regions), showing many filopodia-like processes in the GluD2-OE, but not control, cell. Scale bar, 10 μm. (G) Percentage of control and GluD2-OE cells with 1–2 or 3+ primary dendrites. The numbers of cells within each category are indicated. (H) Percentage of spiny and non-spiny cells according to genotype and number of primary dendrites. The numbers of cells within each category are indicated. (I) Low magnification images of P21 cerebellar cortex showing GluD2 expression levels (I_1_) of cells transfected by IUE that are also labeled by tdT (I_2_). Arrowheads indicate two Purkinje cells with high levels of GluD2 overexpression and shorter dendritic trees. Scale bar, 50 μm. (J) Representative images of a GluD2-OE cell with multiple primary dendrites (arrowheads in J_1_) and high GluD2 expression levels compared to nearby regions (J_2_). Scale bars, 20 μm. (K, L) High magnification confocal images showing dendritic spines in control (K) and GluD2-OE (L) terminal dendritic segments. Scale bar, 5 μm. (M) Number of primary dendrites in control and GluD2-OE cells that have been categorized blindly based on whether they have 1–2 or 3+ primary dendrites. Data are mean ± SEM; the numbers of cells within each category are indicated. (N) Percentage of hyper-spiny and normal cells according to genotype and number of primary dendrites. The numbers of cells within each category are indicated. (O) GluD2 overexpression level according to the number of primary dendrites of GluD2-OE cells with 1– 2 or 3+ primary dendrites. Data are mean ± SEM; the numbers of cells within each category are indicated. (P) Schematic comparing *WT, GluD2* sKO, and GluD2-OE dendritic trees. See Figure S4 for related data.

To test whether these early phenotypes persist in more mature Purkinje cells, we examined GluD2-OE Purkinje cells at P21. Strikingly, some GluD2-OE Purkinje cells had exuberant dendritic branches in the deep molecular layers but failed to extend branches to the pial surface (**Figure 4I**). These Purkinje cells also had supernumerary primary dendrites (**Figure 4J**). While control Purkinje cells have developed well-spaced dendritic spines by this stage (**Figure 4K**), in GluD2-OE Purkinje cells, individual dendritic protrusions were no longer easily resolved and dendritic trunks appeared thicker, suggesting supernumerary dendritic spines (**Figure 4L**). We took an unbiased approach to quantifying these effects by imaging randomly selected transfected cells and then categorizing them based on their number of primary dendrites and, independently, their spine morphology (normal vs. “hyper-spiny”), blind to genotype and GluD2 level. We found that cells with 3+ primary dendrites were less common at P21 than at P7, but all such cells were GluD2-OE cells (**Figure 4M**). Furthermore, whereas all control cells and 95% of GluD2-OE cells with 1–2 primary dendrites had normal dendritic spines, 88% of GluD2-OE cells with 3+ primary dendrites exhibited the hyper-spiny phenotype (**Figure 4N**). Finally, cells with supernumerary primary dendrite phenotype had higher levels of GluD2 overexpression (**Figure 4O**; see **Figure S4A** for representative images of Purkinje cells and their GluD2 intensities). Examination of GluD2-OE cells from unselected samples and additional samples selected based on their shorter dendritic trees (those that failed to reach the pial surface) revealed that Purkinje cells with shorter dendritic trees were associated with higher levels of GluD2 overexpression, supernumerary primary dendrites, and increased thickness along the axis orthogonal to the plane of Purkinje cell dendrite elaboration (**Figure S4B–D**).

Taken together, these data and analyses indicate that GluD2 overexpression causes morphological phenotypes opposite to *GluD2* sparse knockout (**Figure 4P**): over-branching in the deep molecular layer and, in extreme cases, failure to extend dendritic branches to the superficial molecular layer altogether.

### GluD2 That Does Not Bind Cbln1 Cannot Rescue the *GluD2* sKO Phenotypes

If the dendrite morphogenesis defects caused by *GluD2* sKO result from its disruption of parallel fiber→Purkinje cell synaptogenesis, then GluD2’s interaction with its synaptogenic ligand Cbln1 should be required for proper dendrite morphogenesis. To test this prediction, we established an *in vivo* structurefunction assay. The overexpression studies described above indicated that the gain-of-function phenotypes of GluD2 are expression level-dependent (**Figure 4D**), as low levels of IUE-based overexpression of GluD2 did not cause overt dendrite morphogenesis phenotypes. Thus, we used mild expression of wildtype GluD2 to rescue *GluD2* sKO phenotypes by co-electroporating a plasmid encoding wild-type GluD2 (**Figure 5A**). We then compared this to the effect of mild expression of a mutant GluD2 containing four point mutations (D24A, I26A, E61A, R345A; GluD2^DIER^ in short; **Figure 5B**) that abolish binding to Cbln1 (Elegheert et al., 2016). Specifically, we chose cells with somatic GluD2 levels of 25–75% of the neighboring wild-type Purkinje cells, resulting in comparable GluD2 levels between the two conditions (**Figure S5A–C**).

**Figure 5.**
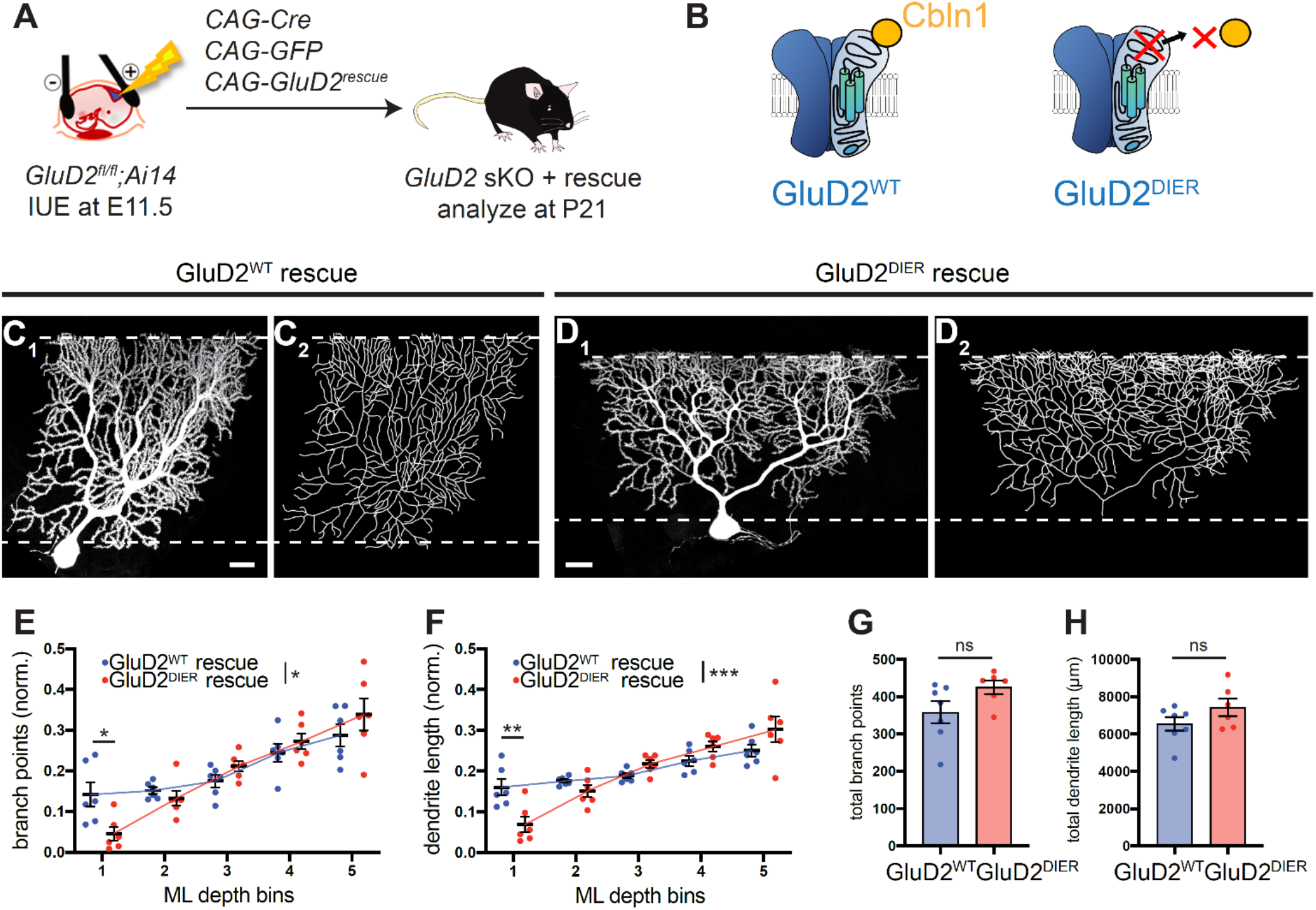
GluD2 that Does Not Bind Cbln1 Cannot Rescue the *GluD2* sKO Phenotypes. (A) Schematic of *in utero* electroporation (IUE) for *GluD2* sKO + GluD2 rescue assay in Purkinje cells of *Ai14* and *GluD2^fl/fl^;Ai14* embryos. Plasmids encoding Cre recombinase, GFP and GluD2^WT^ or GluD2^DIER^ were co-injected into the fourth ventricle at embryonic day 11.5 (E11.5). Cerebellar samples were collected at postnatal day 21 (P21). (B) Schematic illustrating wild-type (left) and mutant “DIER” (right) versions of GluD2. GluD2^DIER^ contains four point mutations (“DIER”, see text) that abolish GluD2 binding to cerebellin-1 (Cbln1), the secreted presynaptic ligand of GluD2. (C) Confocal image (C_1_) and tracing (C_2_) of a representative *GluD2* sKO Purkinje cell expressing a *GluD2^WT^* rescue construct which exhibits similar levels of dendritic elaboration in the deep and superficial molecular layer. Scale bar, 20 μm. (D) Confocal image (D_1_) and tracing (D_2_) of a representative *GluD2* sKO Purkinje cell expressing a *GluD2^DIER^* rescue construct which exhibits decreased branching in the deep molecular layer. Scale bar, 20 μm. (E, F) Quantification of the normalized number of branch points (E) and dendrite length (F) in each molecular layer (ML) bin in GluD2^WT^ rescue (blue) and GluD2^DIER^ rescue (red) Purkinje cells. Data are mean ± SEM; n = 6 (GluD2^WT^ rescue), 6 (GluD2^DIER^ rescue) cells from 2 animals each; p-values were calculated using a two-way ANOVA test (between genotype) followed by Sidak’s multiple comparisons tests between GluD2^WT^ rescue and GluD2^DIER^ rescue values in each bin. (G, H) Quantification of total number of branch points (G) and dendrite length (H) of *GluD2^WT^* (blue) and *GluD2^DIER^* (red) rescue Purkinje cells. Data are mean ± SEM; n = 6 (GluD2^WT^ rescue), 6 (GluD2^DIER^ rescue) cells from 2 animals each; p-values were calculated using unpaired t tests. See Figure S5 for related data.

Compared to GluD2^WT^ rescue Purkinje cells (**Figure 5C**, **Figure S5D**), GluD2^DIER^ rescue Purkinje cells (**Figure 5D, Figure S5E**) exhibited significantly fewer dendritic branches and reduced dendrite length in the deep molecular layer (quantified in **Figure 5E, F**). The total branch points and dendritic length did not differ significantly between these two conditions (**Figure 5G, H**). These results support the notion that dendrite branching defects caused by *GluD2* sKO result from a disruption of signaling mediated by Cbln1-GluD2 interactions.

### *GluD2* Sparse Knockout Phenotypes Are Suppressed by Loss of *Cbln1*

So far, we have shown that sparse but not global knockout of *GluD2* causes marked changes in Purkinje cell dendrite morphology, and that Cbln1-GluD2 binding is required for GluD2’s function in this process. Taken together, these findings suggest that competition between neighboring Purkinje cells for synaptogenesis with parallel fibers, mediated by Cbln1-GluD2 signaling, regulates dendritic branching. If so, one would expect that a complete loss of Cbln1 would remove such competition, and thereby suppress the *GluD2* sKO phenotypes.

To test this prediction, we combined *Cbln1* knockout with *GluD2* sKO. We developed an IUE- and CRISPR/Cas9-based *GluD2* sKO approach using plasmids expressing Cas9 and sgRNAs against *GluD2*, along with GFP as a marker of electroporated cells (**Figure 6A**; **STAR Methods**). GluD2 staining validated elimination of GluD2 expression in *GluD2* CRISPR-based sparse knockout (*GluD2* csKO) cells, but not in control cells with sgRNAs against *lacZ* (**Figure S6A–D**). When introduced into Purkinje cells in *wild-type* animals, *GluD2* csKO caused phenotypes (**Figure 6B, C, S6E, S6F**) similar to those produced by IUE of Cre into *GluD2^fl/fl^* (**Figure 1C, D**) or MADM (**Figure 1I, J**). Quantifications revealed both fewer dendritic branch points and reduced dendrite length in the deep molecular layer and increased dendritic branch points and length in the superficial layer compared to control cells (**Figure 6F, G**). However, *GluD2* csKO in Purkinje cells of *Cbln1^−/−^* mice no longer exhibited the typical sKO dendrite branching phenotypes (**Figure 6D, E, S6G, S6H**, quantified in **Figure 6H, I**). Quantification across the entire dendritic tree revealed that loss of *Cbln1* did not affect total dendritic branch points and length and that *GluD2* csKO reduced dendritic branch points in both *wild-type* and *Cbln1^−/−^* backgrounds (**Figure 6J, K**). Taken together, these experiments demonstrate that the *GluD2* sKO phenotype requires the presence of Cbln1 and support the notion that Cbln1/GluD2-mediated competitive synaptogenesis between neighboring Purkinje cells underlies the dendrite morphogenesis defects exhibited in *GluD2* sKO cells.

**Figure 6.**
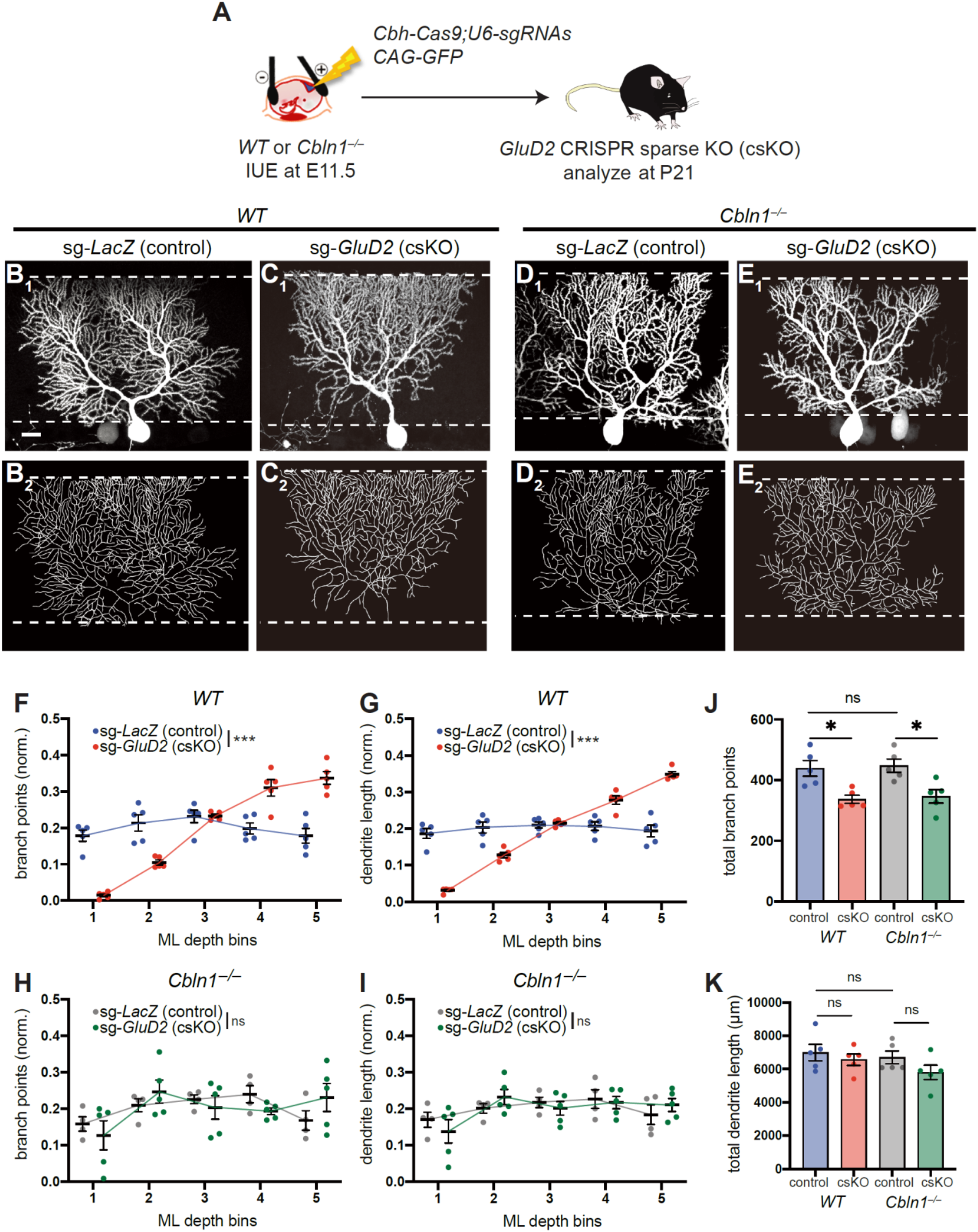
Sparse *GluD2* Knockout Phenotypes Are Blocked by Loss of *Cbln1*. (A) Schematic of *in utero* electroporation (IUE) for CRISPR-mediated sparse control (*LacZ*) or *GluD2* sKO in Purkinje cells of wild-type or *Cbln1^−/−^* embryos. Plasmids encoding GFP and Cas9 (driven by the *Cbh* and *CAG* ubiquitous promoters, respectively), and sgRNAs against *LacZ* or *GluD2* (driven by the *U6* promoter) were co-injected into the fourth ventricle at embryonic day 11.5 (E11.5). Cerebellar samples were collected at P21. (B) Confocal image (B_1_) and tracing (B_2_) of a representative wild-type (WT) Purkinje cell expressing Cas9 and sgRNAs against *LacZ* [sg-*LacZ* (control)], which exhibits similar levels of dendritic elaboration in the deep and superficial molecular layer. Scale bar, 20 μm. (C) Confocal image (C_1_) and tracing (C_2_) of a representative wild-type (WT) Purkinje cell expressing Cas9 and sgRNAs against *GluD2* [sg-*GluD2* (csKO)], which exhibits reduced dendritic elaboration in the deep molecular layer and enhanced dendritic elaboration in the superficial molecular layer. Scale bar, 20 μm. (D, E) Purkinje cells expressing Cas9 and sgRNAs against *LacZ* (D) or *GluD2* (E) in *Cbln1^−/−^* animals both do not exhibit the *GluD2* sKO phenotype, each with similar levels of dendritic elaboration in the deep and superficial molecular layer. Scale bar, 20 μm. (F–I) Quantification of the normalized number of branch points (F, H) and dendrite length (G, I) of Purkinje cell dendrites in each molecular layer (ML) bin in control (sg-*LacZ*: *WT*, blue; *Cbln1^−/−^*, grey) and *GluD2* sKO (sg-*GluD2*: *WT*, red; *Cbln1^−/−^*, green) in wild-type (F, G) or *Cbln1^−/−^* (H, I) animals. Data are mean ± SEM; n = 5 (sg-*LacZ*/*WT*), 5 (sg-*GluD2*/*WT*), 5 (sg-*LacZ*/*Cbln1^−/−^*), 5 (sg-*GluD2*/*Cbln1^−/−^*) cells from 2 animals each; p-values were calculated using two-way ANOVA between sg-*LacZ* and sg-*GluD2* values in each bin; ***, p < 0.001; ns, not significant. (J, K) Quantification of total number of branch points (J) and dendrite length (K) of sg-*LacZ*/*WT*, sg-*GluD2*/*WT*, sg-*LacZ*/*Cbln1^−/−^* and sg-*GluD2*/*Cbln1^−/−^* cells. Data are mean ± SEM; n = 5 (sg-*LacZ*/*WT*), 5 (sg-*GluD2*/*WT*), 5 (sg-*LacZ*/*Cbln1^−/−^*), 5 (sg-*GluD2*/*Cbln1^−/−^*) cells from 2 animals each; p-values were calculated using a one-way ANOVA test followed by Sidak’s multiple comparisons tests between the three indicated columns; *, p < 0.05; ns, not significant. See Figure S6 for related data.

### A Generative Model Recapitulates Key Aspects of Purkinje Cell Dendrite Morphogenesis in Wild-Type and *GluD2* Knockout Conditions

To better understand the dynamics of dendritic growth as well as the consequences of *GluD2* deletion, we developed a generative model of Purkinje cell dendrite morphogenesis. Due to the competitive nature of *GluD2*’s influence on dendrite growth, we modeled the growth of three adjacent wild-type Purkinje cells. This model uses nodes on a 2D lattice to represent synapses with parallel fibers, where occupied nodes indicate an existing synapse and unoccupied nodes indicate the potential for a synapse at that location (**Figure 7A**). In the model, elongation and interstitial branching result from distinct events. Elongation extends an existing branch, while branching generates a new dendritic process that forks off of an existing branch. At every point in time and at each location in the dendritic arbor, the probabilities of elongation and branching are determined by three factors: (1) cell-autonomous drives to continue elongating and branching that lessen as the total number of synapses on the tree increases; (2) repulsion from other dendritic processes in the vicinity of the node (**Figure 7B**); and (3) a force that pulls new dendritic growth upward toward the pial surface. The model iterates through each eligible node in the tree and makes the decision to elongate, branch, or neither based on these probabilities (**Figure 7C**). New parallel fibers progressively enter the simulation over time to model the developmental trajectory of the molecular layer (see **STAR Methods** for details).

**Figure 7.**
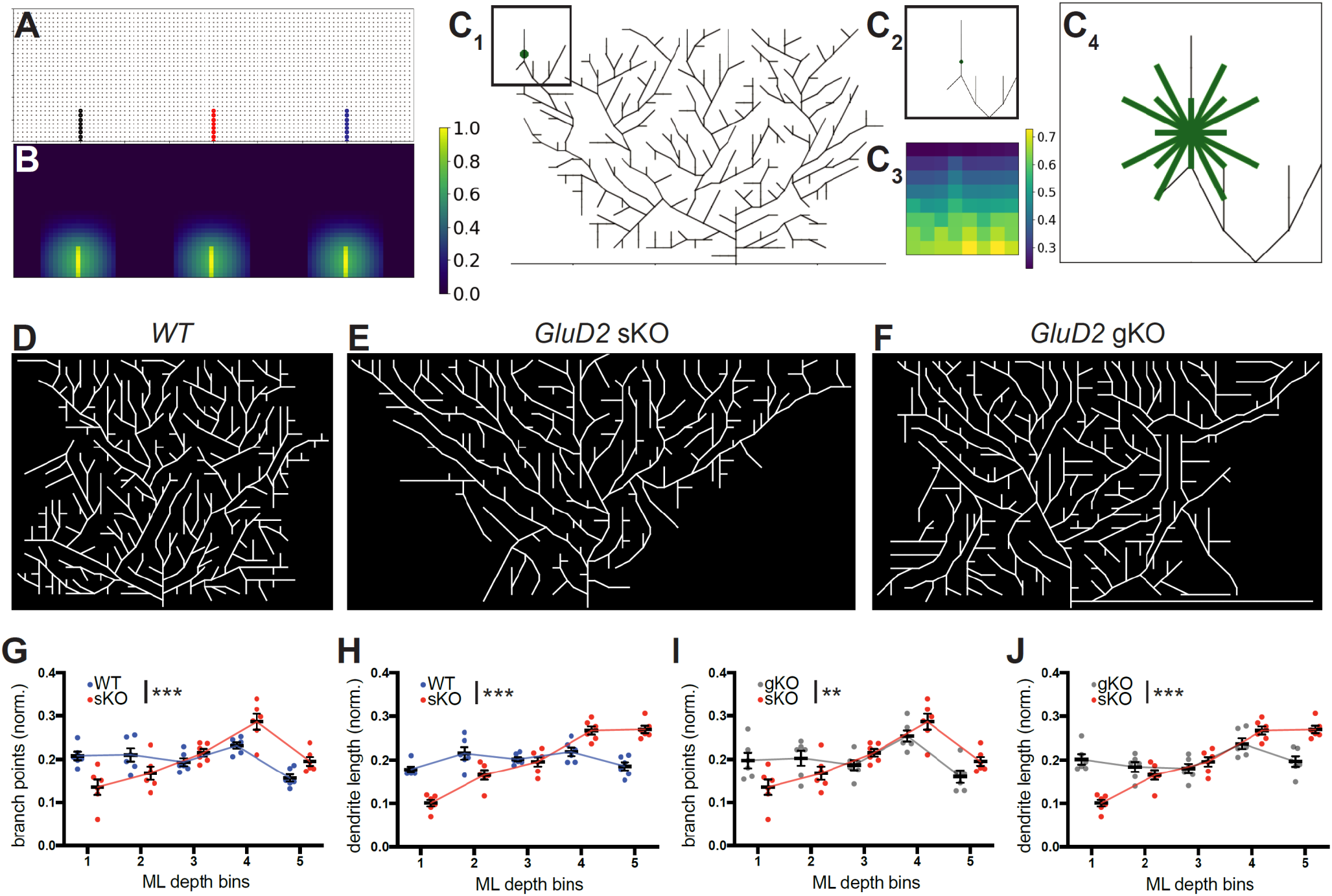
A Generative Model Recapitulates Key Aspects of Purkinje Cell Dendrite Morphology in Wild Type as well as Sparse and Global *GluD2* Knockout Conditions. (A) Skeleton of trees that the model begins with. Each node, denoted by a circle and representing a parallel fiber, is marked occupied when the tree extends a branch through that point on the 2D lattice. This branch (edge) is indicated by a solid line, and the color of the line indicates which tree the edge belongs to (black for the left-most tree, red for the middle tree, and blue for the right-most tree). Nodes with these colors are occupied by the corresponding trees, and other potential nodes for synapses are marked gray. Occupied nodes contribute to repulsion felt in that part of the grid (STAR Methods). (B) Heat map of repulsion throughout the 2D grid shown in (A) with brighter colors indicating stronger repulsion, which discourages growth. For a given cell, the repulsion from nodes occupied by neighboring cells are stronger than repulsion from nodes occupied by itself. Color bar indicates amount of repulsion normalized to the largest value of repulsion in this heat map. (C) Example of a decision a cell might make about whether to branch from a certain point, with eligible directions of branching indicated. (C_1_) A Purkinje cell with a decision point for branching denoted by the green circle. Color bar indicates amount of repulsion normalized to largest value of repulsion in this heat map. (C_2_) The vicinity of the green node shown in (C_1_). (C_3_) As in (B), this is a heat map of the repulsion around the green node, which determines the probability of branching in each direction. (C_4_) Eligible directions for branching from the central node marked by green edges. Note that some of these directions will have 0 probability of branching if they result in crossings with current branches, but all are pictured. The endpoint for each direction is determined by the closest lattice point in that direction; because this introduces different lengths, the probabilities of branching in longer directions are adjusted accordingly. See STAR Methods for detailed description. (D) Representative example of a simulation with three wild-type Purkinje cells, showing the middle cell only (see Figure S7A for the full set of simulations). (E) Representative example of a simulation with one *GluD2* knockout Purkinje cell and two neighboring wild-type cells on either side, showing the middle cell only (see Figure S7B for the full set of simulations). This mimics the sparse knockout experimental condition. (F) Representative example of a simulation with three *GluD2* knockout Purkinje cells, showing the middle cell only (see Figure S7C for the full set of simulations). This mimics the global knockout experimental condition. (G, H) Mean (± SEM) normalized dendritic branch points (G) and length (I) of bins in the molecular layer for wild-type (blue) and sparse *GluD2* knockout (red) conditions. (I, J) Mean (± SEM) normalized dendritic branch points (I) and length (J) of bins in the molecular layer for sparse (red) and global (gray) *GluD2* knockout conditions. For panels G–H, data are mean ± SEM; n = 6 for all three conditions; p-values were calculated using twoway ANOVA between WT/gKO and sKO values in each bin; **, p<0.01; ***, p < 0.001. See Figure S7 for related data, and Movies S1–S3 for illustrations of dendritic growth of three Purkinje cells in WT, sKO, and gKO conditions.

Wild-type cells grown as described tile the 2D grid and display a distinctly square-like morphology (**Figure 7D**, **S7A; Movie S1**). To simulate *GluD2* knockout, we reduced the ratio of that cell’s drive to branch relative to its drive to elongate. This simulates decreased efficacy in stabilizing new synaptic partners and is based on our findings of reduced branching in the initial stages of growth in the deep molecular layer (**Figure 1F**, **3D–F**). We also modified the growth parameters to mimic the reduction of synaptogenesis in *GluD2* knockout cells compared with wild-type cells (Ichikawa et al., 2016). We simulated the sparse knockout condition by only “mutating” the middle cell and observed an inverted triangular morphology like that of sKO cells (**Figures 7E**, **S7B; Movie S2**). Quantification showed fewer dendritic branch points and reduced length in the deep molecular layers (**Figure 7G, H**). The total dendritic length in the superficial molecular layer was also increased in simulated sKO cells compared to controls (**Figure 7H**), but the over-branching phenotype in the superficial molecular layer was less pronounced (**Figure 7G**). This difference with experimental data is likely due to our simulated dendritic trees growing exclusively in a single plane, whereas *GluD2*-deficient cells exhibited increased branching in and out of the dendritic plane (Kaneko et al., 2011), likely accounting for the dendritic branch crossings in the superficial layers of sKO cells (**Figure 1F**; **Figure 3F_3_, inset**). Interestingly, when we mutated all three cells in the 2D grid to simulate gKO conditions, all cells maintained a square-like morphology like that of wild-type cells (**Figure 7F, S7C; Movie S3**; quantified in **Figure 7I, J**). Thus, these simulations recapitulated key features of sparse and global *GluD2* knockout cells, supporting the competitive nature of *GluD2*’s effect on dendritic morphogenesis.

## DISCUSSION

Here, we explore the relationship between synaptogenesis and dendrite morphogenesis by examining the effects of disrupting GluD2, a postsynaptic receptor required for synapse formation and maintenance, on dendrite morphology of cerebellar Purkinje cells. Several lines of evidence support the notion that the dendrite morphogenesis defects we observed in *GluD2* sparse knockout cells were due to disruption of its established function in synapse formation and maintenance. First, we observed a number of defects in *GluD2* knockout cells consistent with disrupted synaptogenesis: prevalence of thin filopodia-like protrusions in place of more mature dendritic spines at P21 (**Figure S1E, F**) and the invasion climbing fibers onto distal dendritic branches of *GluD2* sKO Purkinje cells (**Figure S1G**), as previously described in global *GluD2* knockout mice (Ichikawa et al., 2002). Second, overexpression of GluD2 in Purkinje cells recruited vGluT1 puncta to dendritic branches and produced spine-like protrusions at P7 (**Figure 4**), consistent with GluD2’s role in promoting synapse formation with parallel fibers. Third, the ability to bind Cbln1, its synaptogenic ligand, was essential for GluD2 to rescue reduced dendritic branching in the deep molecular layer due to sparse *GluD2* knockout (**Figure 5**). Fourth, dendrite morphogenesis phenotypes of *GluD2* sKO cells were suppressed by additional loss of *Cbln1* (**Figure 6**), suggesting that they were caused by neighboring Purkinje cells competing to form synapses with parallel fibers. Lastly, by simulating a cell-autonomous synaptogenesis defect in a competitive dendritic growth model, we recapitulated *GluD2* sparse and global knockout morphological phenotypes (**Figure 7**), suggesting that reducing synaptogenesis in a competitive environment is sufficient to cause the dendrite morphogenesis defects we observed.

Sparse *GluD2* knockout Purkinje cells feature two prominent and countervailing characteristics: under-elaboration of the dendritic tree in the deep molecular layer and overelaboration in the superficial molecular layer (**Figure 1**). Morphological analyses across Purkinje cell postnatal development (**Figure 1**; **Figure 3**) reveal that the under-elaboration in the deep molecular layer coincides with reduced branching in the same locale, suggesting that the under-elaboration may be caused in part by a marked local reduction in dendritic branching. Furthermore, GluD2 overexpression caused supernumerary primary dendrites at P7, some of which persisted to P21 (**Figure 4**), suggesting that parallel fiber→Purkinje cell synaptogenesis may help stabilize dendritic branches. These phenotypes resemble those caused by perturbation of the neurexin/neuroligin synaptic organizers in *Xenopus* tectal neurons, where dendritic filopodia are destabilized by neuroligin disruption and stabilized by neuroligin overexpression (Chen et al., 2010). Indeed, the tripartite neurexin/Cbln1/GluD2 synaptic adhesion complex is considered to function in synaptogenesis in the cerebellar cortex analogously to the neurexin/neuroligin complex elsewhere (Südhof, 2018; Yuzaki, 2018). Thus, just as in the *Xenopu*s study (Chen et al., 2010), a reduction in dendritic branching of *GluD2* sKO dendrites in the deep molecular layer supports the synaptotrophic hypothesis (Cline and Haas, 2008; Vaughn, 1989) in a mammalian circuit *in vivo*. We note, however, that whereas both global and sparse perturbations of neuroligin affect dendritic filopodia stability in *Xenopus* tectal neurons (Chen et al., 2010), the reduction of Purkinje cell dendritic branching in the deep molecular layer only occurs in the sparse *GluD2* knockout condition, highlighting the importance of competitive synaptogenesis in the regulation of Purkinje cell dendrite morphogenesis.

The overelaboration of Purkinje cell dendrites in the superficial molecular layer in sparse *GluD2* knockout appears to oppose the prediction of the synaptotrophic hypothesis and highlights a bimodal relationship between synaptogenesis and dendritic growth. We propose two explanations from different perspectives for this overelaboration phenotype. From a holistic perspective, each Purkinje cell’s dendrite growth may be under homeostatic regulation of total synapses formed with parallel fibers (which account for the vast majority of input synapses onto Purkinje cells). Under-elaboration of *GluD2* sKO dendrites in the deep molecular layer may be compensated for by overelaboration in the superficial molecular layer in order to reach a set point of total synapses formed. From a cell biological perspective, the formation of a stable synapse may provide a dendritic growth cone a signal to stop further exploration at later stages of dendrite development. By reducing the ability of Purkinje cell dendrites to form stable synapses with parallel fibers, *GluD2* sKO may thus also prevent cessation of growth, causing exuberant branching and dendritic over-extension. This is supported by our observation of preferential enrichment of long terminal segments in P63 mice (**Figure 3I–J**). We note that these two explanations are not mutually exclusive; indeed, synapse formation as a signal to stop further dendritic exploration could be one of the mechanisms underlying homeostatic regulation. Our generative model contains elements of both perspectives: it features the assumption that an increase in the number of synapses in a dendritic tree results in a homeostatic decrease in the drive for a tree to branch, like a signal to reduce further exploration, and such modelling yielded results similar to experimental observations (**Figure 7**).

A striking finding of this study is that both the under-elaboration and overelaboration of Purkinje cell dendrites in the deep and superficial molecular layer, respectively, were observed only via sparse but not global knockout of *GluD2*. These findings resemble those of our previous study of neurotrophin-3 (NT3)/TrkC signaling, in which sparse but not global knockout of the neurotrophin receptor *TrkC* caused a marked reduction in Purkinje cell dendrite length and branching (Joo et al., 2014). TrkC has been implicated as a synaptogenic receptor in hippocampal and cortical neurons (Takahashi et al., 2011); however, several lines of evidence suggest that the effect of sparse TrkC knockout in dendrite morphogenesis is distinct from its function as a synaptogenic receptor. The synaptogenic role of TrkC is independent of its NT3-binding and kinase activities (Takahashi et al., 2011), whereas TrkC’s role in regulating Purkinje cell dendrite morphogenesis requires its kinase activity and its interaction with NT3 (Joo et al., 2014). Furthermore, unlike *GluD2* sparse knockout, *TrkC* sparse knockout causes a global decrease in dendritic branching and length; indeed, dendritic trees of most *TrkC* sparse knockout Purkinje cells do not extend to the pial surface. In addition, if the homeostasis hypothesis we proposed above were true, loss of NT3/TrkC may disrupt such homeostatic regulation, whereas loss of Cbln1/GluD2 may selectively impact the morphological mechanisms downstream of competitive synaptogenesis without perturbing homeostatic mechanisms. While further exploration of the relationship between NT3/TrkC and Cbln1/GluD2 signaling will enrich our understanding of the mechanisms of dendrite morphogenesis, both studies highlight the competitive nature of dendrite morphogenesis in mammalian central nervous system neurons and reinforce the importance of studying dendrite development using mosaic methods such as *in utero* electroporation and MADM.

## ACKNOWLEDGEMENTS

We thank M. Mishina for *GluD2^fl^* frozen embryos, T.C. Südhof and J.I. Morgan for *Cbln1^fl^* mice, L. Anderson for help in generating the MADM alleles, W. Joo for a previously unpublished construct, M. Yuzaki, K. Shen, J. Ding, and members of the Luo lab, including J.M. Kebschull, H. Li, J. Li, T. Li, C.M. McLaughlin, D. Pederick, J. Ren, D.C. Wang and C. Xu for discussions and critiques of the manuscript, and M. Yuzaki for supporting Y.H.T. during the final phase of this project. Y.H.T. was supported by a JSPS fellowship; S.A.S. was supported by a Stanford Graduate Fellowship and an NSF Predoctoral Fellowship; L.J. is supported by a Stanford Graduate Fellowship; and an NSF Predoctoral Fellowship M.J.W. is supported by a Burroughs Wellcome Fund CASI Award. This work was supported by an NIH grant (R01-NS050538) to L.L.; the European Research Council (ERC) under the European Union’s Horizon 2020 research and innovations programme (No. 725780 LinPro) to S.H.; and Simons and James S. McDonnell Foundations and an NSF CAREER award to S.G.; L.L. is an HHMI investigator.

## AUTHOR CONTRIBUTIONS

Y.H.T., S.A.S., and L.L. designed the experiments; Y.H.T. carried out IUE experiments with help from D.J.L. and S.A.S.; S.A.S. made the DNA constructs; Y.H.T. and S.A.S. performed histology and data analyses with assistance from M.H.; L.J. performed the simulation with guidance from M.J.W. and S.G.; T.R., X.C., and S.H. provided unpublished MADM mice; Y.H.T., S.A.S, L.J., S.G., and L.L. wrote the manuscript with feedback from all authors; L.L. supervised the project.

## STAR METHODS

### KEY RESOURCES TABLE

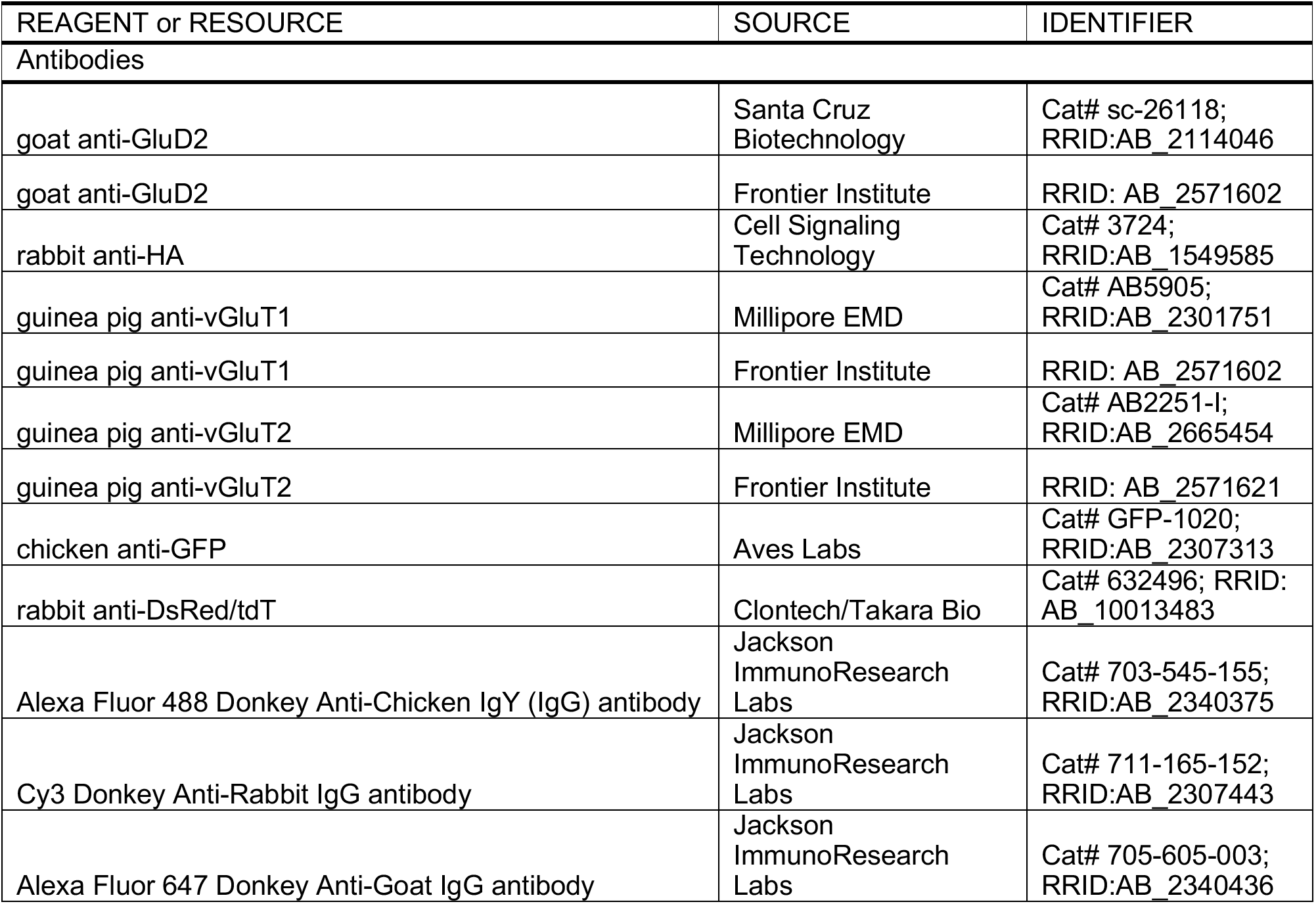
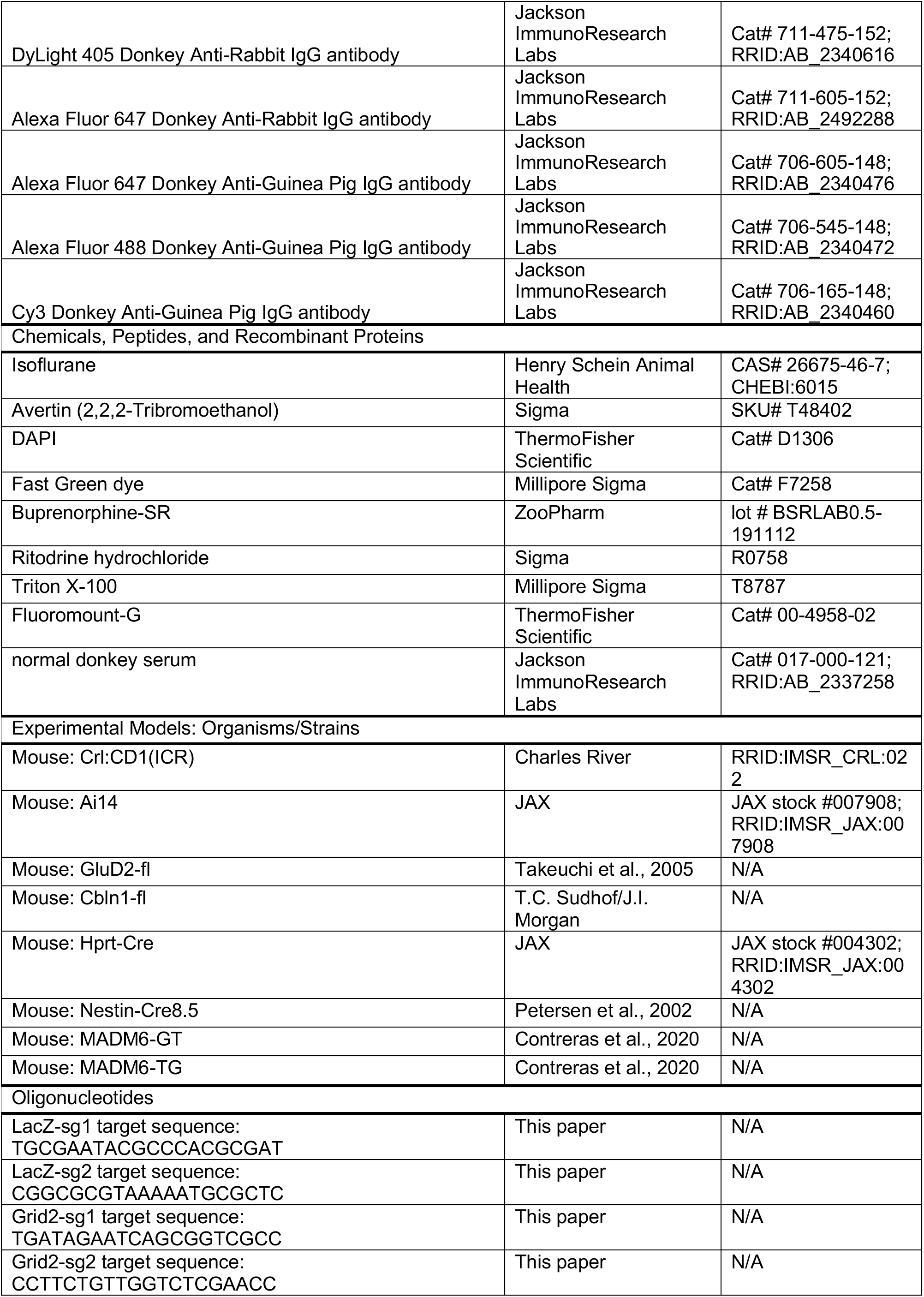
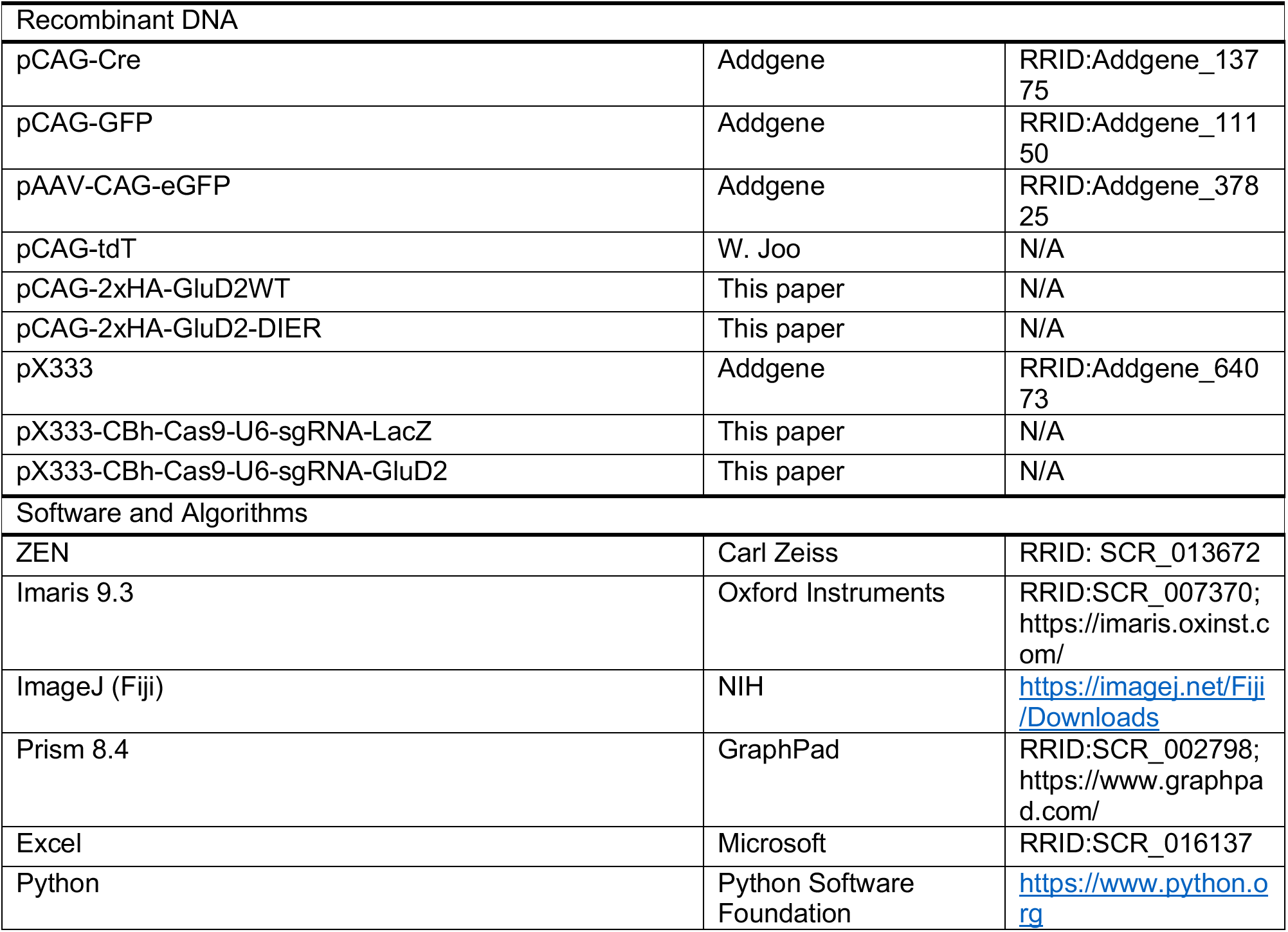

### CONTACT FOR RESOURCE AND REAGENT SHARING

Further information and requests for resources and reagents should be directed to and will be fulfilled by the Lead Contact, Liqun Luo

### EXPERIMENTAL MODEL AND SUBJECT DETAILS

#### Mice

All procedures followed animal care and biosafety guidelines approved by Stanford University’s Administrative Panel on Laboratory Animal Care and Administrative Panel of Biosafety in accordance with NIH guidelines. Mice were housed in plastic cages with disposable bedding on a 12 hours light/dark cycle with food and water available *ad libitum*. Pregnant CD1 dams were ordered from Charles River. *GluD2^fl^* frozen embryos were a kind gift from M. Mishina. *Cbln1^fl^* mice were a kind gift from T.C. Südhof and were generated by J. Morgan. *GluD2^fl^* mice and *Cbln1^fl^* mice were crossed to *Hprt^Cre^* mice obtained from The Jackson Laboratory to generate *GluD2^−/−^* (GluD2 gKO) and *Cbln1^−/−^* mice, respectively. *Ai14* mice were obtained from The Jackson Laboratory. *Nestin-Cre8.5* mice were a kind gift from W. Zhong. MADM-mediated GluD2 sparse knockout mice were generated using MADM6^GT^ and MADM6^TG^ alleles (Contreras et al., 2020) assembled into MADM6-GluD2^fl^ mice as previously described (Joo et al., 2014), with the Nestin-Cre8.5 transgene driving Cre expression in neural progenitor cells.

### METHOD DETAILS

#### *In utero* electroporation

*In utero* electroporation into mouse cerebellar Purkinje cells was performed as described (Nishiyama et al., 2012) with some modifications. Embryonic day 11.5 (E11.5) pregnant dams were anesthetized by isoflurane (starting at 2.5% and maintained at 1.5% in 1 L O_2_/min). After cleaning the abdomen with betadine, a laparotomy was performed, uterine horns were exposed, and DNA was injected within the following 20–30 minutes. To relax the myometrium, ritodrine hydrochloride (0.4–0.8 μg/g; Sigma-Aldrich, St Louis, MO, USA) was injected into the abdominal cavity or directly onto the exposed uterine horns. Warm sterile PBS was continually applied to the embryos to hydrate them. Under the illumination of a fiber optic light source (Dolan Jenner) with a flexible light guide (Allied Electronics), a plasmid DNA solution in a glass capillary needle was injected into the fourth ventricle using a microinjector (Eppendorf FemtoJet 4I; Eppendorf) until the rostral region of the fourth ventricle was filled with DNA, as visualized with Fast Green dye (Sigma). The volume injected into each embryo was approximately 2–3 μL. After injection, the embryo was held through the uterus with tweezer-style electrodes (CUY650P3; NEPAGENE) so that the positive metal electrode was placed on the rostral rhombic lip of the fourth ventricle, and 1–2 sets of electrical pulses (33–38 V, each with a duration of 30 ms, five pulses at intervals of 970 ms per pulse) were delivered using an electroporater (ECM 399, BTX). After electroporation, the uterus was repositioned in the abdominal cavity, 0.05–0.10 mg/kg buprenorphine-SR was injected directly into the intraperitoneal space. The abdominal wall and skin were then sutured closed. The dams were kept on a heating pad until recovery from anesthesia, then returned to their home cages. The embryos were allowed to continue developing and were typically born on E19. After birth, pups were screened for successful electroporation by examining their cerebella through the skin and skull under a fluorescence stereomicroscope, then returned to their home cage with the dam.

#### Cloning and plasmids

Standard cloning procedures were used to generate new DNA constructs. *GluD2* overexpression constructs had two hemagglutinin (HA) tags inserted immediately following the *GluD2* signal sequence and followed by GSG linkers. *GluD2* expression constructs were subcloned into a pCAG vector derived from *pCAG-Cre* (Addgene). The DIER mutant had four point mutations (D24➔A, I24➔A, E61➔A, R345➔A; Elegheert et al., 2016) that abolish binding to cerebellin-1 (Cbln1). Two independent sgRNAs were subcloned into the pX333 plasmid vector (Addgene) for CRISPR-mediated experiments (*sgLacZ* primer sequences: tgcgaatacgcccacgcgat, cggcgcgtaaaaatgcgctc; *sgGluD2* primer sequences: tgatagaatcagcggtcgcc, ccttctgttggtctcgaacc).

Plasmid DNA for IUE was purified using the Qiagen plasmid maxiprep kit (Qiagen) and, following ethanol precipitation, dissolved in HEPES-buffered saline. The plasmid solutions were colored with 0.01% Fast Green so that they were visible when injected into the fourth ventricle. The plasmid DNA used for IUE and their final concentrations were as follows: 1 and 2 μg/μL for *pCAG-eGFP* (or *pCAG-GFP*) and *pCAG-Cre*, respectively (**Figure 1** and **3**); 3 μg/μL for *pCAG-eGFP* or *pCAG-tdTomato* (*pCAG-tdT*) alone (**Figure 2 and 4**); 1 and 2 μg/μL for *pCAG-tdT* and *pCAG-HA-GluD2^WT^*, respectively (**Figure 4**); and 1, 2 and 1 μg/μL for *pCAG-eGFP, pCAG-Cre* and *pCAG-HA-GluD2^WT^*/*pCAG-HA-GluD2^DIER^*, respectively (**Figure 5**); and 1 and 2 μg/μL for *pCAG-eGFP* and *pX333-CBh-Cas9-U6-sgRNA-LacZ/GluD2*, respectively (**Figure 6**).

#### Histology and image acquisition

Mice were deeply anesthetized using 2.5% Avertin and perfused transcardially using 4% paraformaldehyde (PFA) in PBS. The fixed brains were dissected out and postfixed for 1–4 hours at room temperature or overnight at 4°C in 4% PFA in PBS. After washing in PBS, 100-μm thick sagittal cerebellar sections were collected from the cerebellar vermis using a vibratome (Leica), blocked in 10% normal donkey serum in 0.1% Triton X-100 in PBS, and incubated with primary antibodies overnight at room temperature or over two nights at 4°C. Sections were then washed in PBS three times, incubated with Alexa Fluor secondary antibodies (Jackson ImmunoResearch Laboratories) for at least 1 hour at room temperature, washed three times in PBS and mounted and coverslipped on glass slides using Fluoromount-G (Thermo Fisher Scientific). Sections were imaged using a Zeiss LSM 780 laser-scanning confocal microscope (Carl Zeiss).

The z-stacked images for dendrite tracing were acquired using a 40x/1.4 Plan-Apochromat oil immersion objective (Carl Zeiss), at 2048 x 2048 pixels per frame with 0.4 μm z-steps. For IUE-based experiments, Purkinje cells in the bank region of the cerebellar primary fissure were imaged. For experiments using MADM mice, Purkinje cells in the bank regions of the cerebellar primary fissure and lobules III and IV were imaged.

To measure GluD2, vGluT1, and vGluT2 expression and image dendritic spines used in the unbiased analyses in **Figure 4**, an 20X/0.8 Plan-Apochromat air immersion objective (Carl Zeiss) and an 40X/1.4 Plan-Apochromat oil immersion objective (Carl Zeiss) were used. Images in **Figure 4I** were collected using a 10X/0.3 Plan-Neofluar air immersion objective (Carl Zeiss). For certain experiments, P7 Purkinje cells from lobules III and IV/V (Figure 4B–H) and P21 Purkinje cells from lobules III–VIII were imaged (Figure 4I–O).

For **Figure S1E**, z-stack images of dendritic spines were acquired using an 63X/1.4 Plan-Apochromat oil immersion objective (Carl Zeiss) at 2048 x 2048 pixels per frame, with a zoom factor of 3 and 0.4 μm z-steps. For each Purkinje cell, two z-stack images of isolated distal dendrites were obtained, one from the deepest molecular layer depth bin and another from the most superficial bin. Each image included 1 or 2 distal dendrite segments.

#### Antibodies

We used the following primary antibodies: goat anti-GluD2 (Santa Cruz Biotechnology or Frontier Institute; 1:200), rabbit anti-HA (Cell Signaling Technology; 1:500), guinea pig anti-vGluT1 (Millipore Sigma or Frontier Institute, 1:200), guinea pig anti-vGluT2 (Millipore Sigma, or Frontier Institute, 1:200). In some MADM experiments, chicken anti-GFP (Aves; 1:500), rabbit anti-DsRed (Clontech; 1:500) were used.

#### Image analysis and processing

Imaris 9.3 FilamentTracer (Oxford Instruments) was used to trace the dendrites of cerebellar Purkinje cells from z-stack confocal images (see above). Dendrites were traced using semi-automatic AutoPath and Manual modes with a fixed filament diameter of 5 pixels. The dendrite beginning point was defined as the location where the primary dendrite thickness is 8 μm in diameter for P14 and older Purkinje cells. For Purkinje cells at P7 and P10, because the primary dendrites tend to be thicker than that of older ages (see **Figure 3C_1_** and **C_4_**), the thickness at the dendrite beginning point was defined as 10 μm in diameter. All dendritic protrusions longer than 2.5 μm were traced as dendrites. Only three dendritic segments were allowed to form a single dendrite branch point. After tracing all the dendrites, total dendritic length, total number of branch points and total number of dendritic segments were automatically computed by the software and obtained via the Statistics function.

Quantification of dendrite branch points and dendrite length in the molecular layer depth bins were performed as follows. To determine the heights of molecular layer depth bins for each Purkinje cell, two points were marked in an orthogonal view to the x-y plane at the superficial surface of the molecular layer on each side of the Purkinje cell where no labeled dendrites existed (the molecular layer surface was visible by background fluorescent signal). A straight line connecting the two points was drawn and defined to be the superficial surface of the molecular layer. Another straight line passing through the dendrite beginning point was drawn parallel to the superficial surface of the molecular layer. When the Purkinje cell had multiple primary dendrites, the dendrite beginning point of the longest dendrite was used. The region between those two lines was defined as the molecular layer, and the distance between the lines was divided equally into five sublayers, which constitute the molecular layer depth bins. The branch points or dendrites above the 5^th^ (most superficial) bin and those below the 1^st^ (deepest) bin were included into the 5^th^ and 1^st^ bins, respectively. To quantify the dendrite branch points in each bin, x-y coordinates of all branching points were obtained via the FilamentTracer Statistics function. The x-y coordinates were rotated and translated so that the molecular layer was parallel to the x-axis. The dendrite branch points in each bin were sorted according to their y coordinates. To quantify dendrite lengths, the traced dendrites (“filament” objects) were converted into a z-stack image of a dendritic skeleton using the Create Channel function of Imaris9.3 XTension. The dendritic skeleton had a uniform dendritic thickness (5 pixel). The images were opened using ImageJ (or Fiji, NIH), rectangular selections of each bin were made using the ROI Manager, and signal intensities of each ROI were measured from all of the z-sections and summed. The resulting signal intensities were divided by the total signal intensities to determine the relative dendrite lengths in each bin.

Quantification of dendritic spine head widths was performed using ImageJ (NIH). For each spine oriented into the x-y plane, the largest diameter of the spine head was drawn with a straight line perpendicular to the dendritic spine shaft and quantified. This quantification was performed blind to the genotypes and molecular layer depth bins.

For quantification of GluD2 levels in **Figure S5C**, GluD2 levels in cytosolic regions of the somata of labeled and neighboring non-transfected control Purkinje cells were quantified. Background GluD2 levels were defined as the GluD2 levels in the internal granular layer and were subtracted from the somata measurements. The percentages of resulting GluD2 intensities in the labeled cells to those of nontransfected control Purkinje cell were defined to be the relative GluD2 levels. To quantify normalized GluD2 or vGluT1 levels in dendrites at P7 (**Figure 4**), GluD2-expressing dendritic regions were selected using Otsu’s thresholding and the mean intensities of GluD2 and vGluT1 were measured from a labeled cell and its neighboring non-transfected dendrites. Background intensities were defined as the darkest regions in the external granule layer and subtracted from the selected regions. The resulting values of the labeled dendrites were divided by those of non-transfected dendrites and normalized to average GluD2 or vGluT1 values determined in the same manner from control Purkinje cell dendrites expressing only tdT. To quantify normalized GluD2 levels in dendrites at P21 (**Figure S4**), tdT^+^ dendrite regions were selected using Otsu’s thresholding and GluD2 levels were defined to be the GluD2 levels in the labeled cells. As control GluD2 levels, neighboring unlabeled molecular layer regions were selected using the Rectangular tool. Background intensities were defined as the signal in neighboring interneurons’ nuclei and subtracted from the selected regions. The percentage of GluD2 levels of labeled dendrites to that of non-transfected regions was normalized to that of control Purkinje cell dendrites only expressing tdT.

The Purkinje cells in **Figure S4C_2_** and **D**, due to their increased dendrite tree thickness, often had other labeled cells in their close vicinity. Therefore, to highlight the dendrite morphology of these cells, masked images were created. To do so, dendrites belonging to other cells were removed from each z-section using ImageJ (NIH). The modified images were then opened in Imaris9.3 and Surface objects were created based on the modified images. The original raw images were then masked using the Surface objects.

### QUANTIFICATION AND STATISTICAL ANALYSIS

Prism 8.4 (GraphPad) and Excel (Microsoft) were used for data analysis and plotting. All statistical tests were performed using Prism 8.4 (GraphPad).

### MODELLING AND SIMULATION METHODS

#### Overview of basic algorithm

We simulated Purkinje cell dendritic tree growth on a single 2D plane pierced by orthogonal intersecting parallel fibers (PFs). Specifically, we started with 3 adjacent Purkinje cells (PCs) at an early developmental stage (**Figure 7A**). Growth occurs either through extension of dendritic segments from their terminals (elongation) or interstitial branching from an existing non-terminal segment (which we hereafter refer to as branching). Both elongation and branching result in the growth of a new segment. Every dendritic segment consists of a straight line that starts and ends at nearby discretized lattice points on the 2D parallel fiber grid. Each such lattice point corresponds to a potential PF→PC synapse, which is then realized when a dendritic segment occupies that lattice point. Note that in our simplified 2D planar growth model, we did not consider influences from PCs at other 2D planes. We modelled only PF→PC synapses, which accounts for a vast majority of synapses onto PCs (tens of thousands per PC), and omit the impact of climbing fiber inputs (several hundred per PC).

The dendritic segments of a modeled PC can be oriented along one of 16 directions, corresponding to 16 possible nearby lattice displacements between the start and end of every dendritic segment: (*Δx*, *Δy*) ∈ {(0, ±1), (±1,0), (±1, ±1), (±1, ±2), (±2, ±1)} (Figure 7C_4_). These 16 choices of lattice displacements correspond to an approximate angular separation of 22.5 degrees between adjacent directions. Our final lattice grid of PFs extends across 3 × 38 = 114 lattice points horizontally and 38 lattice points vertically. We model 3 adjacent PCs under 3 different conditions: (1) WT, (2) *GluD2* sparse knockout (sKO) (i.e., modifying growth parameters of the central PC only), or (3) global knockout (gKO) (changing the growth parameters of all three PCs). The probability of successful elongation or branching at any given point to one of the 16 directions follows the general rules below for all PCs, albeit with different growth parameters for WT and *GluD2* knockout cells:

1. To restrict excessive dendrite turning, elongation from a terminal point is limited to 3 directions (0 degrees or approximately ± 22.5 degrees) relative to the orientation of the terminal segment from which elongation is initiated.
2. Branching can occur in any one of the 16 directions.
3. The probability of elongation and branching upwards is higher (mimicking a presumptive attractant from the pial surface or a repellent from internal granular layer).
4. To enforce lack of collisions, neither elongation nor branching can result in the creation of a new dendritic segment that terminates at a lattice point already occupied by a previously grown dendritic segment, and no new segment can intersect any such previous segment.
5. In addition, both the probabilities of elongation and branching to regions with a high density of nearby dendritic segments are diminished through specific modelling of longer range repulsion: dendritic self-avoidance from the same tree and tiling of dendritic trees of neighboring cells.

To execute these rules, at each iteration we randomly picked an occupied lattice point of one of the three existing dendritic trees. Any such chosen lattice point can belong to one of three classes: (1) it is a terminal point on the tree, in which case, in our model, elongation is the only possibility for growth from this point; (2) it is a non-terminal point from which the dendrite extends in two directions, in which case branching is the only possibility for growth from this point; (3) it is a branch point in which the tree already extends in three directions, in which case we do not allow further growth from this point. For the randomly chosen point, our model makes a probabilistic decision as to whether or not to elongate or branch when feasible.

More specifically, let E and B be the events that successful elongation or branching, respectively, actually occurs from a candidate growth point on the tree. The probability *P*(*E*) [or *P*(*B*)] of successful elongation (or branching) from that growth point is not necessarily 1 and could actually fail due to hard constraints involving collision avoidance or soft factors due to longer range repulsion in all possible directions. However, when a successful elongation or branching event actually occurs, the direction *θ* along which growth occurs is also chosen probabilistically from the conditional distributions *P*(*θ*|*E*) for elongation and *P*(*θ*|*B*) for branching. In general, directions *θ* in which appreciable dendrodendritic repulsion is far away or directions that are closer to the upwards direction are favored in the distributions *P*(*θ*|*E*) and *P*(*θ*|*B*). This process of randomly choosing a growth point, randomly deciding whether or not to grow, and then randomly deciding which direction to grow, is repeated for 2000–4000 iterations to grow all three trees.

Below, we describe in sequence how we modeled dendrodentritic repulsion, how this repulsion determines the conditional probability distributions *P*(*θ*|*E*) and *P*(*θ*|*B*) for which direction *θ* to elongate or branch respectively, and how the total level of repulsion, along with intrinsic drives for growth, determine the probabilities *P*(*E*) and *P*(*B*) of successful growth or elongation in the first place. We then describe which growth parameters are different in WT and *GluD2* knockout cells. Finally, we end with a high-level intuitive description of the key principles enabling our model to account for the experimental data.

#### Modeling dendrodentritic repulsion

We define *r_i_*(*x,y*) to be the repulsion field of neuron *i* = 1,2,3, which always equals the convolution of the current dendritic tree with a spatially decaying kernel. The kernel occupies 17 × 17 lattice points with a central value of 1.875 and all other elements of the kernel fall of as 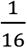. Euclidean Distance from the center. This kernel, when convolved with the dendritic tree of neuron *i*, yields a spatially decaying repulsion field *r_i_*(*x,y*) around each tree. Examples of the three repulsion fields are shown at an early growth state in **Figure 7B**. The growth of a given tree involves a total repulsion field *r*(*x, y*) that combines its own self-repulsion field with that of its neighbors. For example, for the middle cell, *r*(*x, y*) = 5 ∗ *r*_1_(*x, y*) + *r*_2_(*x, y*) + 5 ∗ *r*_3_(*x, y*); this ensures that neighbor repulsion is five times stronger than self-repulsion.

#### Repulsion as a function of angle determines which direction to grow

To determine the conditional probabilities *P*(*θ*|*E*) and *P*(*θ*|*B*) conditioned on a successful elongation (E) or branching (B) event, we first define a growth score *s*(*θ*) that describes how favorable it would be to grow in a direction *θ*, where *θ* is one of the 16 possible directions for growth shown in **Figure 7C_4_**. For growth directions *θ* in which a collision would occur, *s*(*θ*) = 0, which effectively forbids growth. Otherwise, 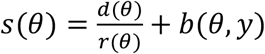. In the first term, *d*(*θ*) is the distance along the growth direction *θ* to L(K) the nearest lattice point (*x, y*) with nonzero repulsion *r*(*x, y*) and *r*(*θ*) is the value of this repulsion. Thus directions *θ* in which appreciable repulsion is far away yield a larger value of the ratio 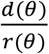, thereby increasing the growth score *s*(*θ*).

The second term *b*(*θ, y*) promotes vertical growth, with increasing strength the closer the growth point is in *y* to the pial surface and the closer the growth angle *θ* is to the upward direction. More specifically, we define 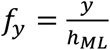 to be the fraction of the vertical distance *y* the growth point is to the height of the molecular layer *h_ML_*. For angles *θ* in which Δ*y* ≤ 0, *b*(*θ, y*) = max(0.75,1 – *f_y_*). For angles *θ* in which (*Δx*, *Δy*) = (±2,1), *b*(*θ,y*) = 0.5 · *f_y_*. For angles *θ* in which (*Δx*, *Δy*) = (±1,1), *b*(*θ,y*) = *f_y_*. For angles *θ* in which (*Δx*, *Δy*) = (±1,2), *b*(*θ,y*) = 1.5 · *f_y_*. For the vertical *θ* = 90 degrees in which (*Δx*, *Δy*) = (0,1), *b*(*θ,y*) = 2 · *f_y_*. Basically, the vertical bias score simply increases linearly with both the height of the growth point and with the rank ordering of the possible slopes. Thus, in summary, the growth score *s*(*θ*) is larger when appreciable repulsion is further away in direction *θ*, and also larger when this direction is closer to upwards. However, it is 0 if growth in direction *θ* would lead to a collision. Finally, this growth score determines the angular conditional probability distributions of growth through

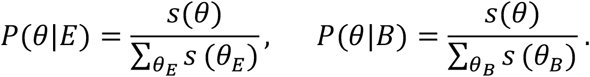

Here, for elongation, the sum over *θ_E_* extends over only 3 angles: 0 degrees or approximately ± 22.5 degrees, relative to the orientation of the terminal segment from which elongation is initiated and *P*(*θ*|*E*) can be nonzero only for these 3 angles, yielding a normalized probability distribution. Conversely, the sum over *θ_B_* extends over all 16 possible directions. In summary, these probabilities determine the randomly chosen angle of growth, by forbidding collisions, avoiding the repulsion field, favoring upwards growth, and restricting dendritic turning for elongation but not branching.

#### A balance between total repulsion and intrinsic drive determines the decision to grow

We next describe how our model computes the probabilities *P*(*E*) and *P*(*B*) of successful elongation or branching events in the first place. Both these probabilities are inversely related to the total repulsion *R*_TOT_ summed across all directions and directly related to overall growth drives *E*(*n*_syn_) and *B*(*n*_syn_). These latter drives depend on the number of synapses *n*_syn_ already formed and stabilized by the tree and are decreasing functions of *n*_syn_, reflecting that the total growth drive decays as the number of synapses increases. Thus, in our model, synapse formation acts as a soft slow-down inhibitor of dendritic growth. Specifically, the probabilities of successful growth are obtained by balancing the growth drives against the total repulsion through

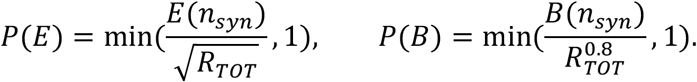

Thus, high total repulsion *R*_TOT_ in the local vicinity of a candidate growth point and larger numbers of stable synapses *n*_syn_ across the entire tree both decrease the probabilities of successful elongation or branching from that growth point. We next describe how we compute the total repulsion *R*_TOT_ and the growth drives *E*(*n*_syk_) and *B*(*n*_syk_).

The total repulsion *R*_TOT_ is defined as follows. Let *R*(*θ*) be a measure of the repulsion in direction *θ*. We define 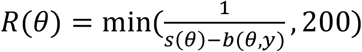 where *s*(*θ*) is the growth score defined above. Basically, in a direction *θ* in which growth would lead to a collision, *R*(*θ*) would take a maximal penalizing value of 200. Conversely, in a direction *θ* in which growth would lead to no such collision, 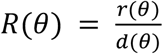, where, again, *d*(*θ*) is the distance along the growth direction *θ* to the nearest lattice point (*x, y*) with nonzero repulsion *r*(*x, y*), and *r*(*θ*) is the value of this repulsion. Then we have *R*_TOT_ = ∑_*θ*_ *max*(1, *R_θ_*)). In simulations, *R*_TOT_ is typically on the order of 10^0^ to 10^1^.

Now the growth drives are given by

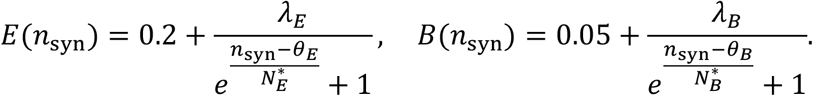

Both are sigmoidal decreasing functions of the number of synapses *n*_syn_ already formed. *θ_E_* and *θ_B_* are threshold parameters; when *n*_syn_ is significantly above either threshold, the corresponding growth drives starts to diminish. Correspondingly, 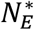 and 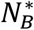 set the scale, in terms of number of synapses above threshold, at which diminished growth starts to set in. Finally, *λ_E_* and *λ_B_* are overall constants that contribute to the corresponding growth drive when *n*_syn_ is at the corresponding threshold *θ_E_* or *θ_B_*.

For wild-type PCs, 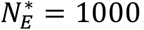, *θ_E_* = 500, and *λ_E_* = 0.5. This yields a modulation of *E*(0) ≈ 0.5 at the beginning of growth and an *E*(1000) ≈ 0.4 at a late stage of growth. Also, for wild-type PCs, 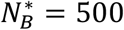, *θ_B_* = 250, and *λ_B_* = 10 for wild-type PCs. This yields a modulation of *B*(0) ≈ 6.3 at the beginning of growth and a *B*(1000) ≈ 1.9 at a late stage of growth. Then when combined with the total repulsion, for wild-type PCs the elongation probability *P*(*E*) is in the range 0.01 to 0.02 in early growth and is close to 0.009 in later growth stages. In contrast, the branching probability *P*(*B*) is typically in the range 0.01 to 0.025 in early growth stages, and close to 0.007 in later stages.

#### Parallel Fiber Expansion

Once the dendritic trees for each condition have elaborated enough such that branches stop growing (due to excessive repulsion in every direction), we expanded the grid of parallel fibers available as synaptic partners. We did so by adding a set of rows and 3 times that number of columns (due to having 3 cells). After the expansion, each cell has an equal amount of excess space to grow both upwards and to either side. The parallel fiber lattice starts as a 90×30 lattice, and then expands to 102×34 and finally to 114×38.

#### Modeling the loss of the GluD2 receptor

To simulate the effect of losing the GluD2 receptor, we made two changes:

6. We change *λ_E_* and *λ_B_* from (*λ_E_, λ_B_*)_*WT*_ = (0.5,10) to (*λ_E_, λ_B_*)_*mut*_ = (4,0.5). Thus, the intrinsic growth drive for elongation is increased, while the growth drive for branching is decreased. We justify this change based on experimental evidence that there were fewer branching points in the 2D plane for knockout cells than for WT cells in the deep molecular layer (**Figure 3**). Also, a decrease in branching likelihood simulates a tree being less effective at growing outward (which involves growing at more angles than just upwards or at a 22.5-degree offset from upward). Because this model is probabilistic and each branch is final, having a decrease in likelihood of branching simulates the process of two growth cones, one from a knockout cell and one from a wild-type cell, competing for a synapse with a PF and the knockout growth cone losing and thus retracting. Then the likelihood of elongation is also increased so as to compensate for the consequent reduction in number of synapses formed—the PC dendrites extend throughout the molecular layer even without the GluD2 receptor; indeed, *GluD2* sKO cells overextended dendrites beyond the upper border of the molecular layer at P10 (**Figure S3J**).
7. The knockout cell is less effective at forming synapses because loss of GluD2 will diminish its ability to stabilize synapses. Experimental evidence also suggests that only 60% of synapses are retained in a *GluD2* knockout condition. Thus, we stochastically count 60% of the number of synapses total in a given knockout cell (used to parametrize the *E*(*n*_syn_) and *B*(*n*_syn_) curves), which is on average equivalent to replacing *n*_syn_ with 0.6 · *n*_syn_ for the knockout cell. This means that the knockout cell will experience a diminished reduction in growth drives *E*(*n*_syn_) and *B*(*n*_syn_).

Overall, this simple change of only two parameters *λ_E_* and *λ_B_* in going from WT to knockout is sufficient to account for two striking experimentally observed morphological phenotypes: square dendritic trees in the WT condition (where all three cells are WT), and an inverted triangular dendritic tree for the middle knockout cell in the sKO condition (with a WT neighbor on either side). While we made various choices about model parameters, we do not claim that these are the only sets of parameters that will result in these phenotypes. This is certainly one such set that mimics the experimental data, but there is likely a diversity of parameter choices that can explain the data.

#### Key principles underlying model behavior

Regardless of detailed parameter choices, a few key principles explain the behavior of the model. First, in going from WT to knockout, elongation becomes favored while branching is suppressed. In the sKO condition, the suppressed branching forces the knockout cell to lose the competition with its WT neighbors to grow its tree by forming synapses in the deep layers where two WT and one sKO tree are all initially present. Thus, the WT neighbor dendritic trees invade the territory of the sKO tree and prevents its outward branching in the deep layers. However, the enhanced elongation of the sKO cell relative to WT cells enables it to reach the superficial layers before the WT does. Moreover, even if the WT tree catches up to the sKO tree in the superficial layers, it has a more diminished drive for branching [reduced *E*(*n*_syn_) and therefore reduced *P*(*B*)] because it has already formed more synapses than the sKO tree due to its already winning the competition in the deep layers (i.e., it has a larger *n*_syn_).

Thus, a combination of three effects enables the sKO cell to win the competition for territory with its WT neighbors in the superficial layers despite the fact that it lost the competition in the deep layers: (1) the sKO tree reaches the superficial layer earlier, (2) the WT cell has diminished drive for branching in the superficial layers due to a larger number of already formed synapses in the deep layers, and (3) the sKO has a less diminished drive for branching in the superficial layers due to formation of fewer synapses in the deep layers. Combined, these effects explain the emergence of the inverted triangular phenotype of the sKO cell through the loss of a competition with its WT neighbors in the deep molecular layer and the win of a competition with its WT neighbors in the superficial molecular layer.

### DATA AVAILABILITY

Data will be made available upon reasonable request to the lead contact.

---

**Figure S1.**
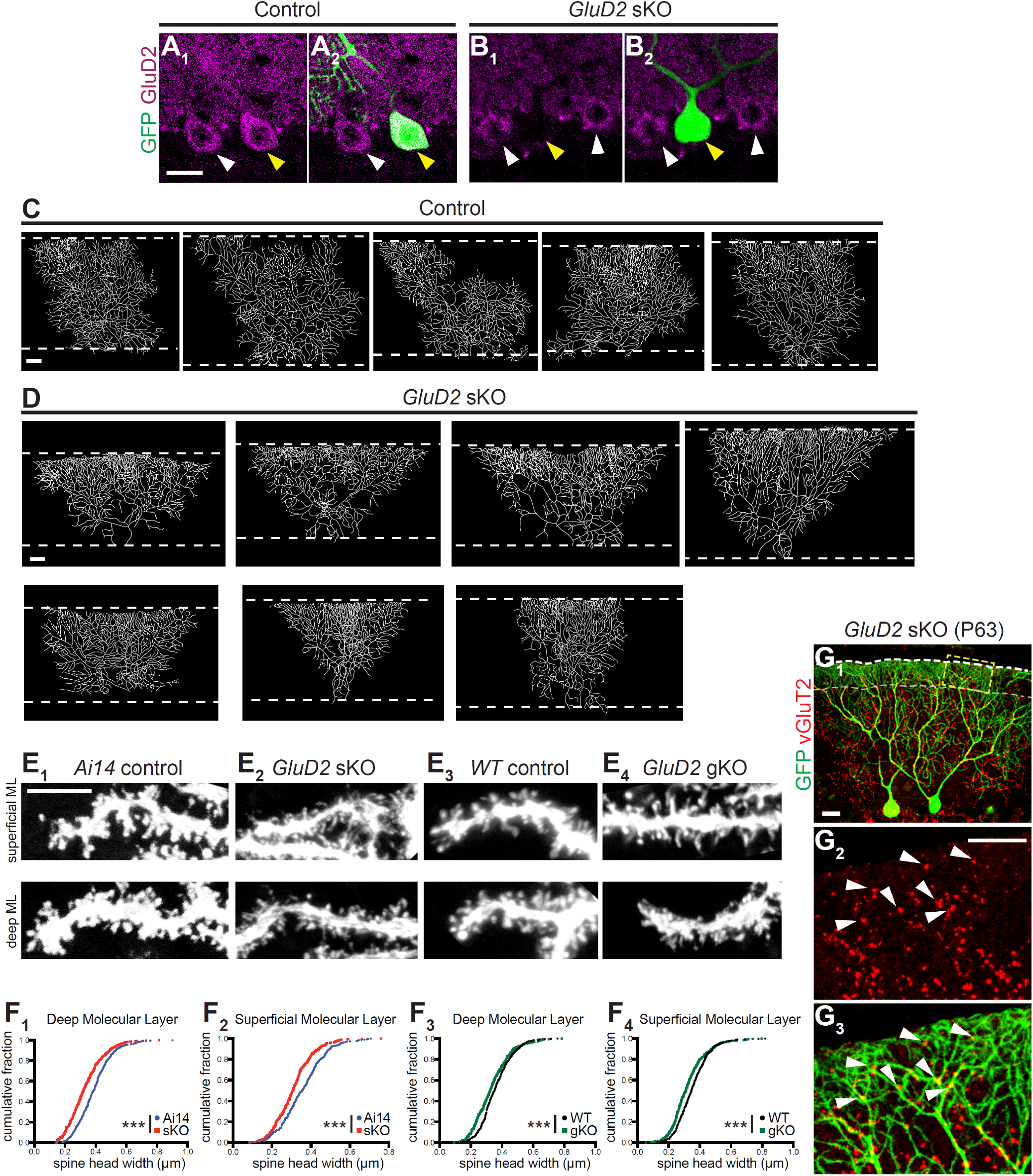
Morphological Characteristics of *GluD2* sKO Purkinje Cells Generated via *In Utero* Electroporation, Related to Figure 1. (A, B) GluD2 staining in representative GFP^+^ transfected control (A) and *GluD2* sKO (B) cells and neighboring cells. Yellow arrowheads indicate normal GluD2 staining in a control cell (A_1_) and a lack of GluD2 staining in a *GluD2* sKO cell (B_1_). White arrowheads indicate unlabeled neighboring cells. Scale bar, 20 μm. (C, D) Dendritic tracings of control (C) and *GluD2* sKO (D) Purkinje cell dendrites. Scale bar, 20 μm. (E) Representative dendritic segments showing spine morphology in *Ai14* control (E_1_), *GluD2* sKO (E_2_), *WT* control (E_3_), and *GluD2* gKO (E_4_) cells at P21. Scale bar, 5 μm. (F) Quantification of dendrite spine head widths of *Ai14* control, *GluD2* sKO, *WT* control and *GluD2* gKO cells. Data show cumulative distributions; n = 444 (*Ai14* control), 442 (*GluD2* sKO), 405 (*WT* control), 414 (*GluD2* gKO) deep molecular layer spines (F_1_, F_3_); and n = 417 (*Ai14* control), 414 (*GluD2* sKO), 383 (*WT* control), 453 (*GluD2* gKO) superficial molecular layer spines (F_2_, F_4_); from 2 (Ai14 control), 3 (*GluD2* sKO), 2 (*WT* control), 2 (*GluD2* gKO) animals; p-values were calculated using the Kolmogorov-Smirnov test; ***, p < 0.001. (G) Representative image of vGluT2 staining in low (G_1_) and high magnification (G_2_, G_3_, for vGluT2 only and vGluT2/GFP, respectively) images, showing expansion of vGluT2 puncta (arrowheads) to the superficial molecular layer apposed to GFP^+^ *GluD2* sKO cells. vGluT2 is a presynaptic terminal marker expressed only by climbing fibers at this stage. The gray dashed line represents the superficial border at which vGluT2 staining normally terminates. Scale bar, 20 μm.

**Figure S2.**
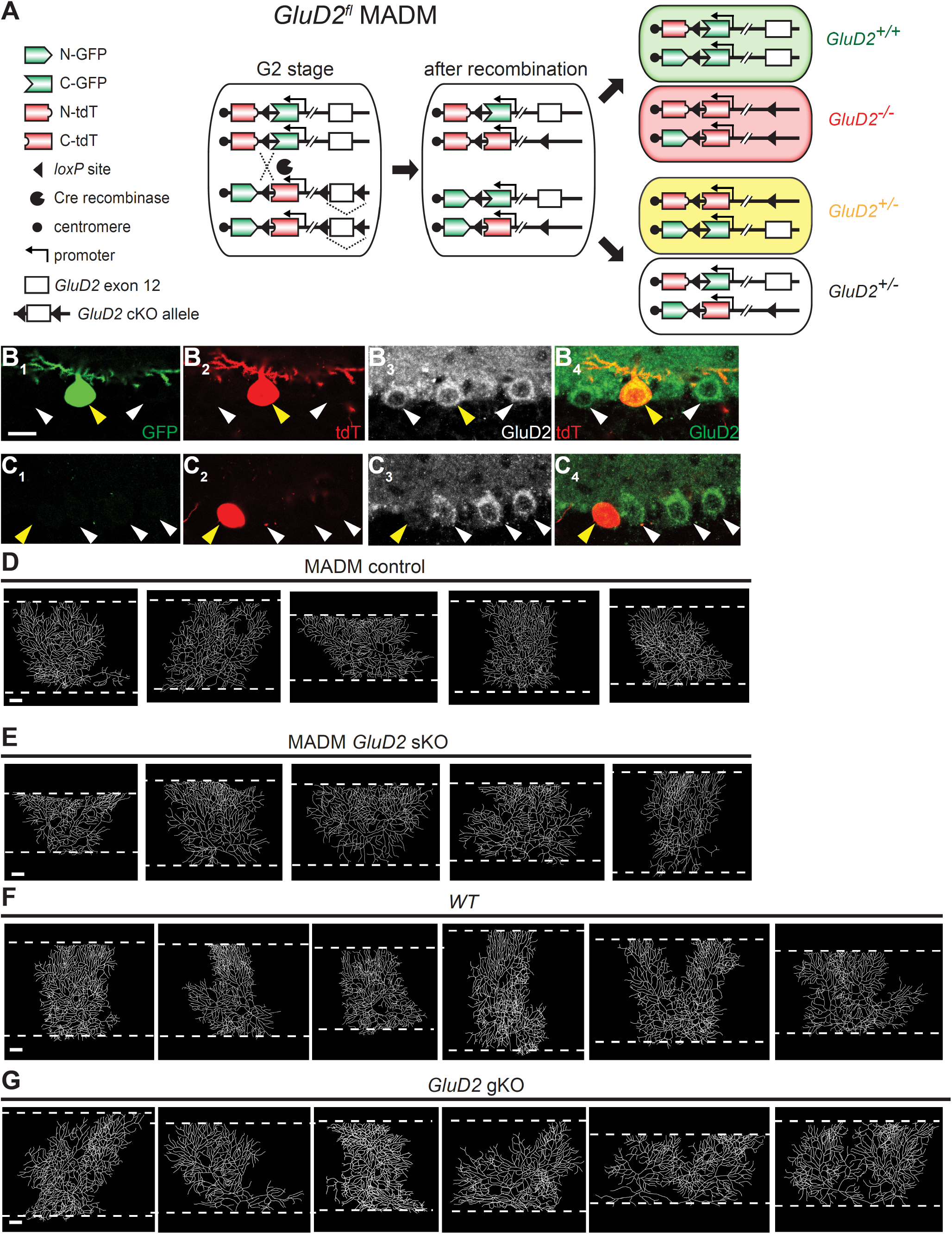
Morphological Characteristics of MADM *GluD2* sKO and Global *GluD2* Knockout Purkinje Cells, Related to Figures 1 and 2. (A) MADM schematic illustrating inter-chromosomal recombination leading to sparsely-labeled *GluD2^+/+^, GluD2^+/−^*, and *GluD2^−/−^* (*GluD2* sKO) cells. See Contreras et al. (2020) for detail. (B, C) GluD2 staining in representative MADM-labeled GFP^+^/tdT^+^ *GluD2^+/−^* control (B) and GFP^−^/tdT^+^ *GluD2^−/−^* sKO (C) cells, as well as neighboring *GluD2^+/−^* cells not labeled by MADM. Yellow arrowheads indicate normal GluD2 staining in a MADM control cell (B_3_) and a lack of GluD2 staining in a MADM *GluD2* sKO cell (C_3_). White arrowheads indicate unlabeled neighboring cells. (D, E) Tracings of MADM control (*GluD2^+/−^*, D) and GluD2 sKO (*GluD2^−/−^*, E) Purkinje cell dendrites. (F, G) Tracings of control (*wild-type*, F) and *GluD2* global knockout (gKO, G) Purkinje cell dendrites, related to Figure 2. Scale bars, 20 μm.

**Figure S3.**
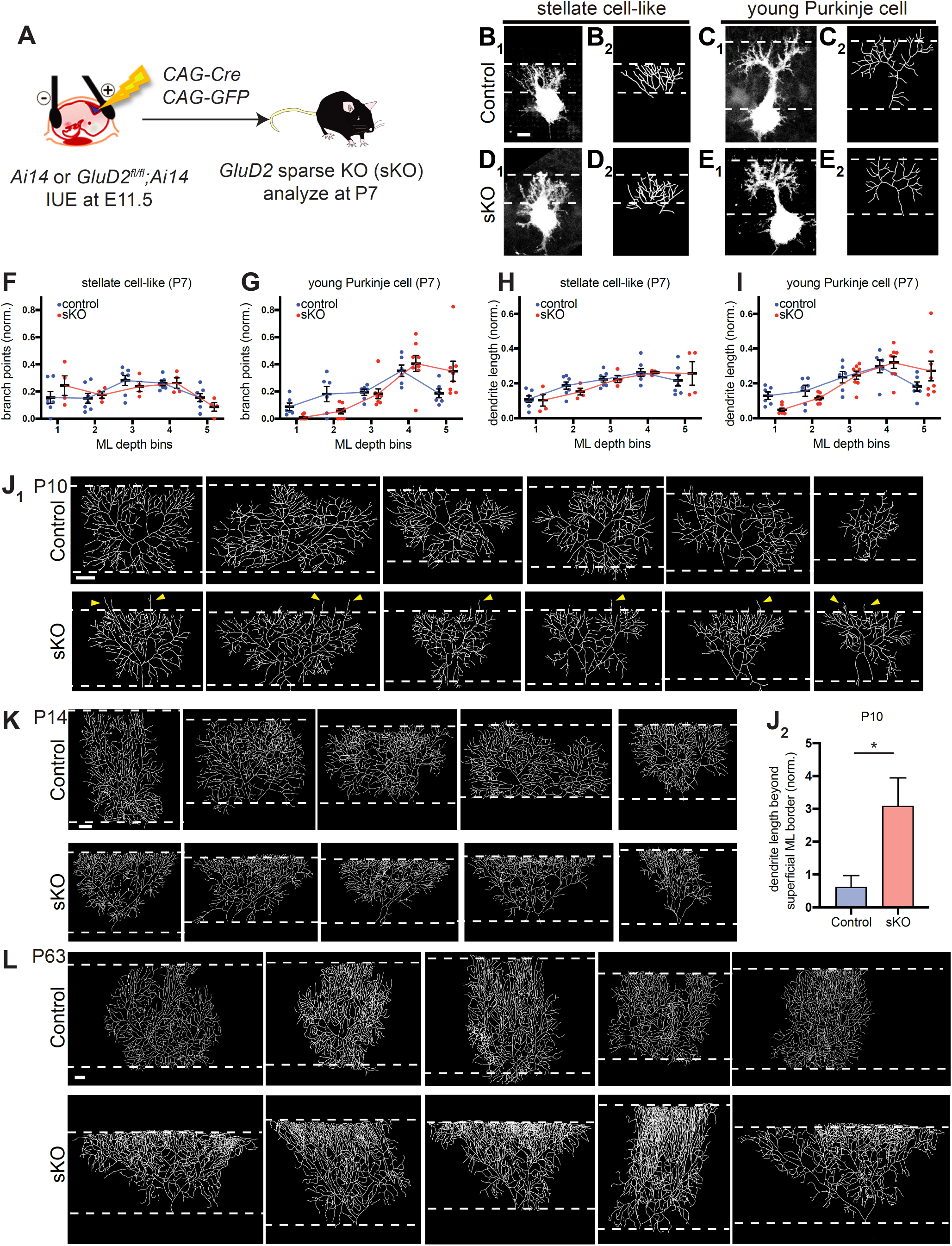
Additional Data on Developmental Analysis of *GluD2* sKO, Related to Figure 3. (A) Schematic of *in utero* electroporation (IUE) for genetically accessing Purkinje cells in *Ai14* and *GluD2^fl/fl^;Ai14* embryos. Plasmids encoding Cre recombinase and GFP were co-injected into the fourth ventricle at embryonic day 11.5 (E11.5). Cerebellar samples were collected at postnatal day 7 (P7). (B–E) Representative confocal images (B_1_–E_1_) and tracings (B_2_–E_2_) of control (B, C) and *GluD2* sKO (D, E) Purkinje cells at P7. These trees were categorized as stellate cell-like (with 3+ primary dendrites) or young Purkinje cells (with 1–2 primary dendrites). Scale bar, 10 μm. (F–I) Quantification of the normalized number of branch points (F, G) and dendrite length (H, I) in each molecular layer (ML) depth bin in control (blue) and *GluD2* sKO (red) Purkinje cells in the stellate cell-like (F, H) and young Purkinje cell categories (G, I). While control and sKO stellate-like cells appear similar at P7, in the young Purkinje cell category, there is a trend toward sKO cells having fewer dendritic branch points and reduced proximal dendrite length in bin 1 and more dendritic branch points and increased distal dendrite length in bin 5. (J–L) Dendritic tree tracings of P10 (J_1_), P14 (K), and P63 (L) control and *GluD2* sKO cells. J_2_, Quantification of normalized dendrite length that extends beyond the superficial border of the molecular layer. Data are mean ± SEM; n = 6 (control), 6 (sKO) cells from 2 animals each; p-value was calculated using a two-tailed, unpaired t test. *, p < 0.05. Scale bars, 20 μm.

**Figure S4.**
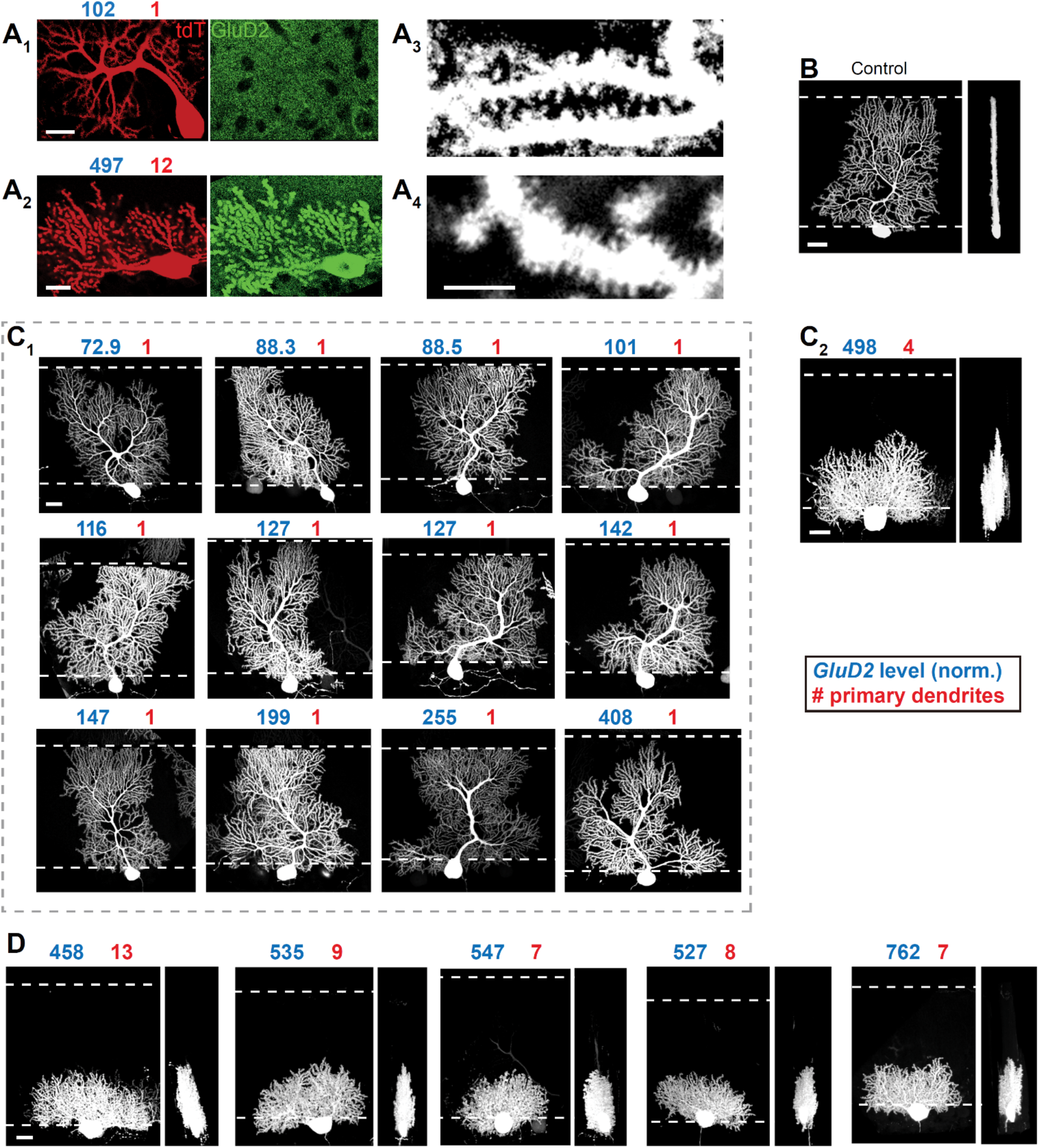
Morphological Characteristics of GluD2-OE Purkinje Cells, related to Figure 4. (A) Representative images of GluD2 staining (A_1_–A_2_) and dendritic spines (A_3_–A_4_) of GluD2-OE Purkinje cells used for quantifications shown in **Figure 4M–O**. GluD2 intensity (relative to control) and number of primary dendrites are indicated with blue and red numbers, respectively. Scale bars, 20 μm (A_1_–A_2_) and 5 μm (A_3_–A_4_). (B) Left, representative z-stack confocal image of a control Purkinje cell at P21. Right, 90-degree rotation of the image, revealing the cell’s planarity. (C) Z-stack confocal images of GluD2-OE Purkinje cells collected in an unbiased manner. Cells were categorized post hoc based on whether they have 1–2 (C_1_) or 3+ (C_2_) primary dendrites. (C_2_) Left, z-stack confocal image of a GluD2-OE Purkinje cell with 4 primary dendrites. Right, 90-degree rotation of the image, revealing a substantially thicker dendritic tree. (D) Unbiased imaging of entire dendritic trees tended to exclude Purkinje cells with short dendritic trees (e.g., C_2_) due to their increased dendritic thickness, which was often cut through in the tissue sections. Therefore, we further selected GluD2-OE Purkinje cells from new animals, based on their short dendritic trees. Post hoc measurements indicated high GluD2 overexpression levels (blue numbers) and supernumerary primary dendrites (red numbers). Right, images after 90-degree rotations revealing increased dendritic tree thicknesses. Scale bars: 20 μm (B–D).

**Figure S5.**
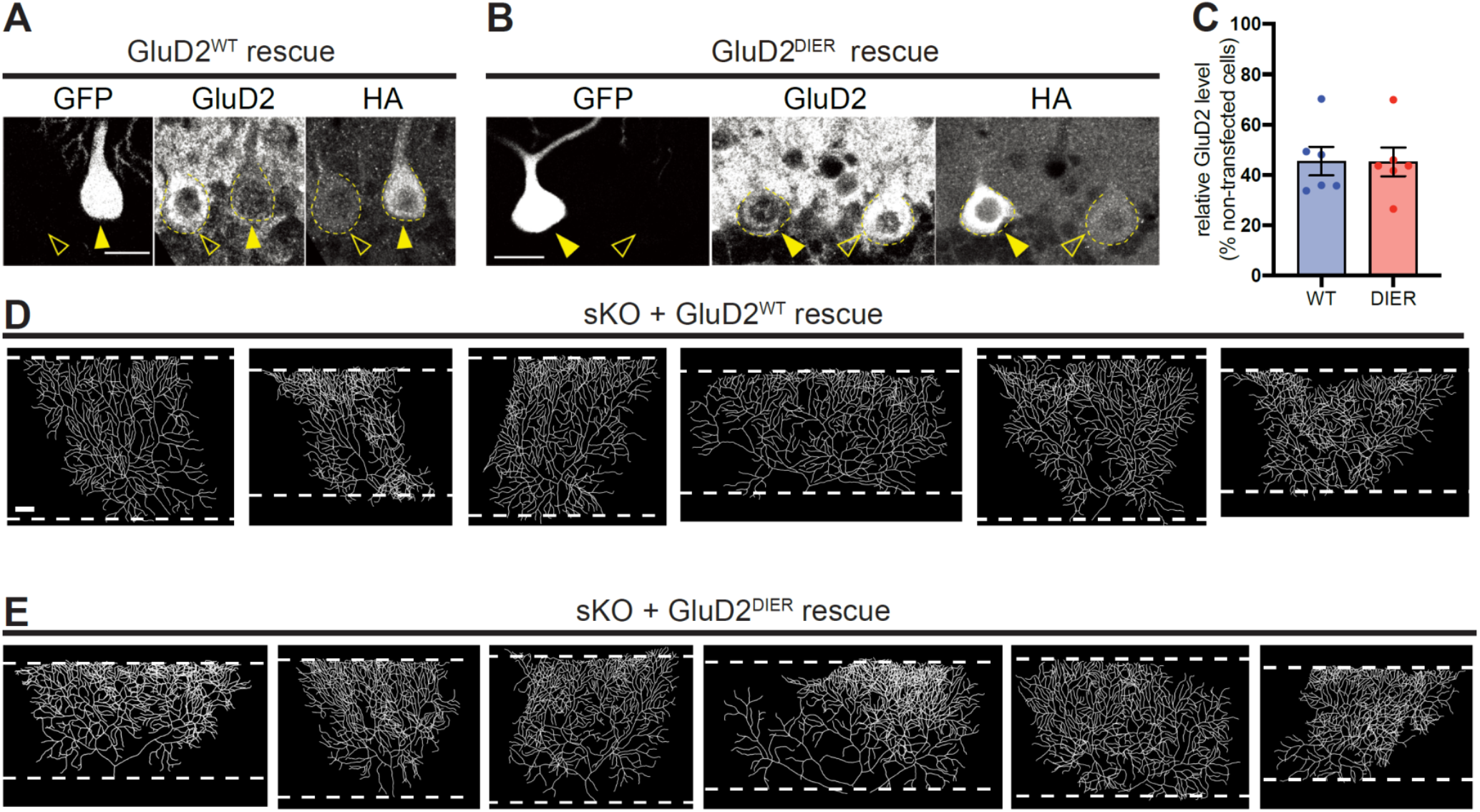
Additional Data on the GluD2 Rescue Experiments, Related to Figure 5. (A, B) Representative images showing GluD2 level in GluD2^WT^ rescue (A) and GluD2^DIER^ rescue (B) cells. Filled arrowheads indicate cells co-transfected with *GluD2* and *GFP* transgenes (as indicated by both HA and GFP staining). Open arrowheads indicate neighboring untransfected *wild-type* cells. (C) Quantification of GluD2 staining levels in GluD2^WT^ rescue (A) and GluD2^DIER^ rescue (B) cells. (D, E) Dendritic tracings of GluD2^WT^ rescue (D) and GluD2^DIER^ rescue (E) cells. Scale bars, 20 μm.

**Figure S6.**
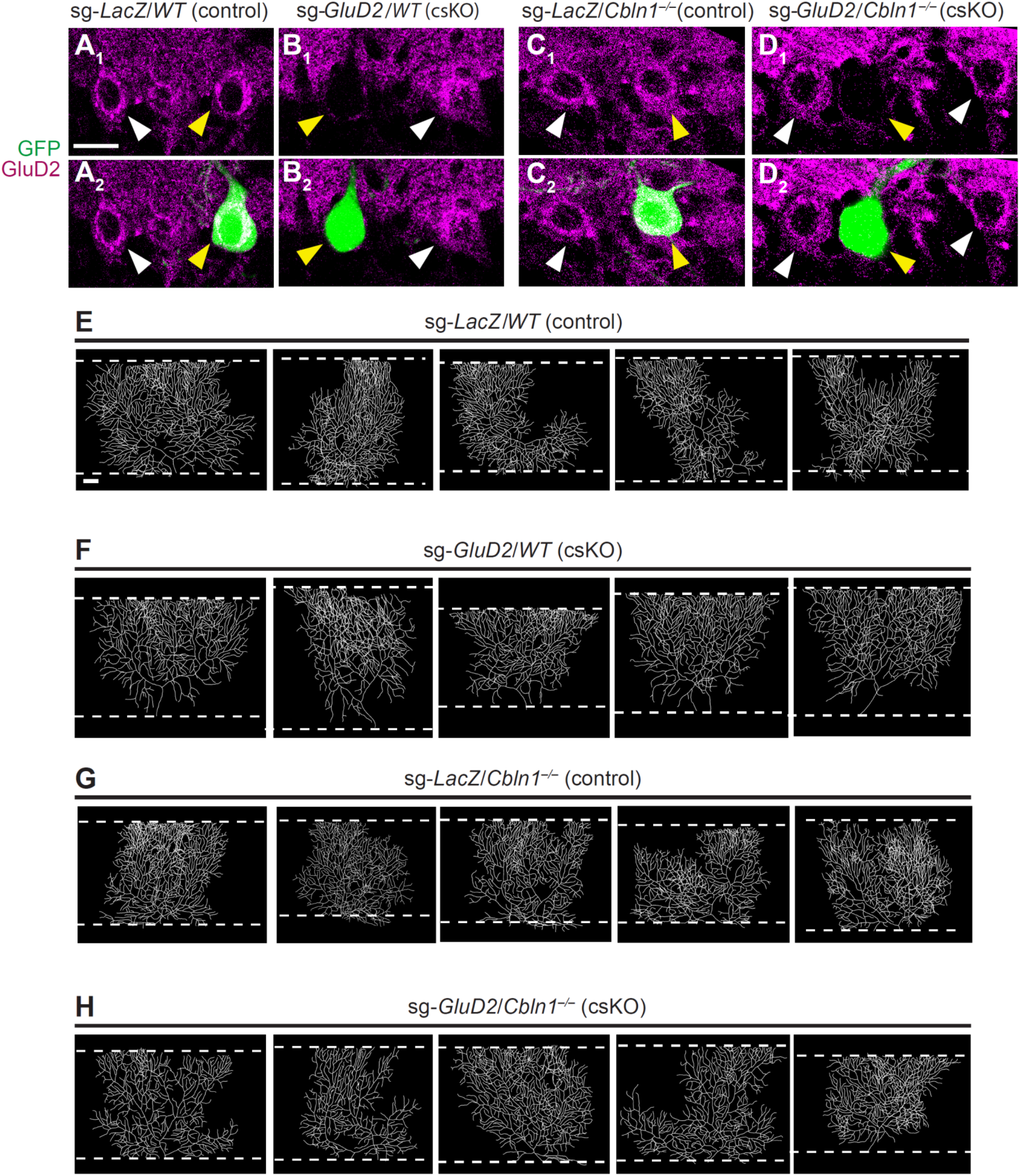
Additional Data on CRISPR-mediated *GluD2* sKO in Wild-type and *Cbln1^−/−^* Backgrounds, Related to Figure 6. (A–D) GluD2 staining of Purkinje cells transfected with sg-*LacZ* (A, C) and sg-*GluD2* (B, D) in *wild-type* (A, B) and *Cbln1^−/−^* (C, D) backgrounds. Yellow arrowheads indicate GFP^+^ transfected cells showing a lack of GluD2 staining in sg-*GluD2* but not sg-*LacZ* conditions, when compared to neighboring nontransfected cells indicated by white arrowheads. (E–H) Dendritic tracings of sg-*LacZ*-transfected *WT* (E) and *Cbln1^−/−^* (F) cells and sg-*GluD2*-transfected *WT* (G) and *Cbln1^−/−^* (H) cells. Scale bar, 20 μm.

**Figure S7.**
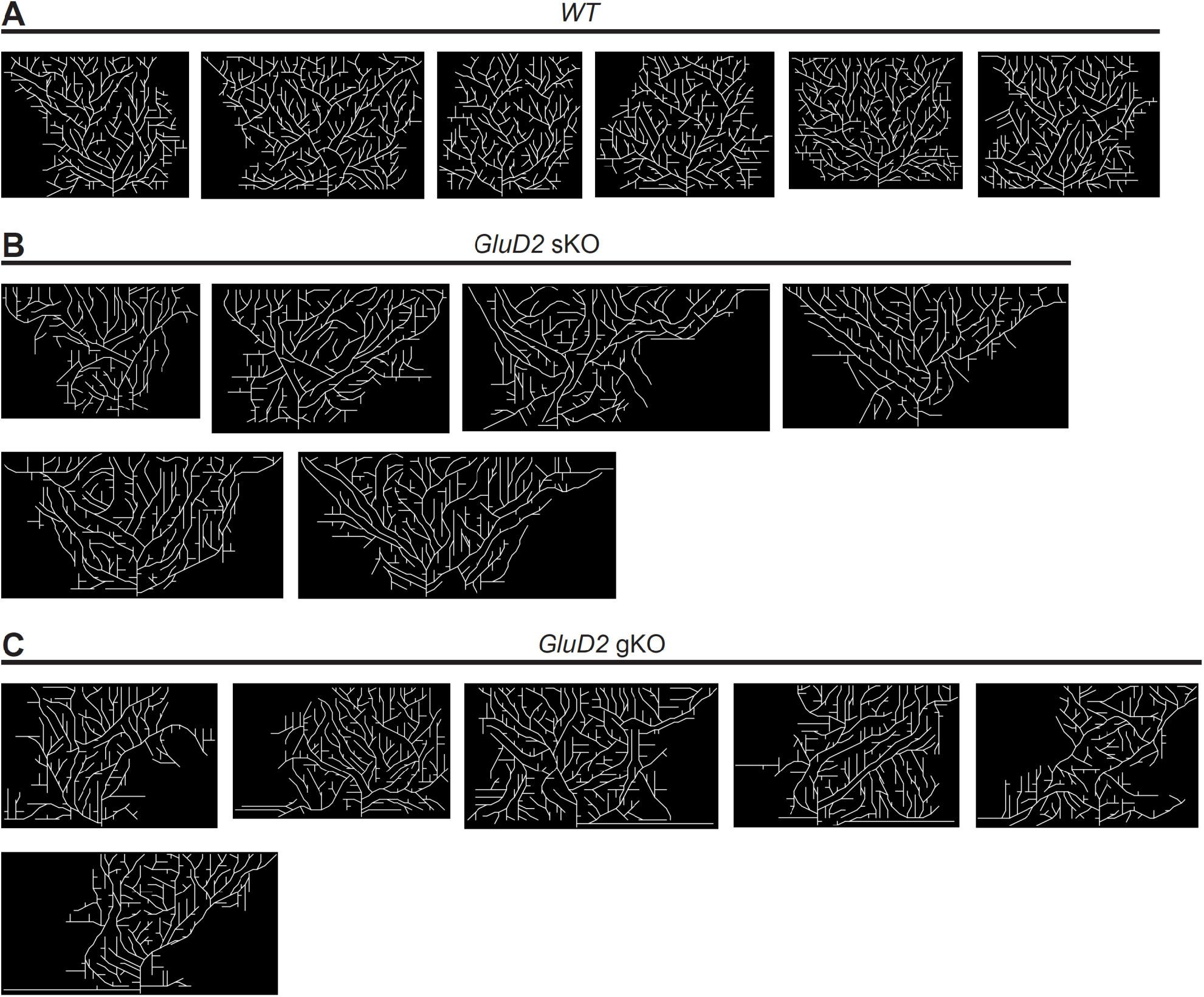
Full Set of Simulations Results, Related to Figure 7. (A) Full set of simulations with three wild-type Purkinje cells, showing the middle wild-type (*WT*) cell only. (B) Full set of simulations with one *GluD2* knockout Purkinje cell and two neighboring wild-type cells on either side, showing the middle cell only. This mimics the sparse knockout (sKO) experimental condition. (C) Full set of simulations with *GluD2* knockout Purkinje cells, showing the middle cell only. This mimics the global knockout (gKO) experimental condition.

**Movie S1. Simulations of Dendritic Growth of Three Wild-type (WT) Cells.**

**Movie S2. Simulations of Dendritic Growth of One *Glud2^−/−^* (KO) Cell Flanked by Two Wild-type (WT) Cells.**

**Movie S3. Simulations of Dendritic Growth of Three *Glud2^−/−^* (KO) Cells.**

